# Mapping Tumor-Microenvironment dependencies with TMEformer: A spatial foundation framework enabling in silico perturbation

**DOI:** 10.64898/2026.05.17.725770

**Authors:** Shensuo Li, Panting Liu, Qiheng Chen, ZiChao Zhang, Yingkui Tang, Dian Liu, Gongsheng Xu, Mingzhu Zhou, Jinqin Luo, Lan Huang, Bogong Chen, Siyang Ou, Jingwen Jiang, Xiyi Wei, Lu Yang, Guanghui Zhu, Sujun Chen

## Abstract

Despite the fundamental role of spatial context in driving tumor progression, most current computational models for virtual perturbation have largely overlooked its importance. Here, we introduce TMEformer, a tumor microenvironment-aware deep learning framework that leverages high-resolution spatial transcriptomics to jointly model intrinsic tumor cell programs and local microenvironmental signals by explicitly incorporating spatial architecture. Validated across diverse tumor spatial transcriptomic cohorts, TMEformer enables virtual perturbations that capture functional dependencies within local cellular ecosystems. Despite being trained on cancer-specific spatial datasets, TMEformer outperforms baseline models pretrained on large-scale corpora in capturing key tumor transitions, including lineage plasticity and the emergence of therapy resistance. Systematic perturbation analyses prioritize tumor-intrinsic transcription factors and TME-derived ligands that drive disease progression, recovering established regulators and revealing novel candidates. Furthermore, TME- derived embeddings improve the spatial stratification of tumor cells and align more closely with pathological architecture. Together, TMEformer establishes a general framework for modeling tumors as spatially coupled, perturbable ecosystems. An online TMEformer web platform for perturbation prediction, analysis, and visualization is available at http://www.pradcellatlas.com.

## Introduction

With the advent of single-cell and spatial omics, tumor development and progression are increasingly recognized as emergent properties of complex cellular ecosystems rather than the consequence of tumor cell–intrinsic alterations alone^1, 2^. Intratumoral heterogeneity, lineage plasticity, and therapy resistance reflect the capacity of malignant cells to dynamically adapt to diverse selective pressures^3^. These adaptive processes cannot be fully understood without considering the spatially organized tumor microenvironment (TME), particularly in solid tumors, which provides biochemical signals, physical constraints, and intercellular interactions that continuously shape tumor cell states^4^. However, analyses for the tumor ecosystem defined by tumor cells and surrounding stromal and immune compartments remain largely descriptive, emphasizing co-localization patterns or static interactions. A central unresolved challenge remains in deciphering how perturbations in tumor-intrinsic or microenvironmental compartments propagate through the spatial network to orchestrate tumor cell states^5, 6^. Addressing this challenge requires computational frameworks that are capable of modeling tumor–TME interactions as integrated systems.

Concurrently, transformer-based foundation models have revolutionized single-cell representation learning^7, 8^. By pretraining on large-scale single-cell omics data, these models extract generalizable gene regulatory patterns, facilitating downstream tasks such as cell type annotation and batch correction. This paradigm has given rise to the concept of the artificial intelligence virtual cell (AIVC)^9^. Leveraging this, in silico perturbations (ISP) enable the simulation of genetic or pharmacological interventions to predict cellular responses, ultimately offering a scalable route to hypothesis generation and mechanistic inference^10^. However, most single-cell models treat cells as isolated entities and often rely on *in vitro* data input ^11^, which may not fully recapitulate the *in vivo* responses shaped by the complex tumor microenvironment (TME) ^12–14^. As spatial transcriptomics (ST) continues to advance,^5^ pioneering works like MintFlow ^15^ and SpaceTravLR ^16^ have begun to bridge this gap. Nevertheless, considerable untapped potential remains in harnessing non-linear deep learning architectures to capture the intricate, non-canonical regulatory logic underlying cellular organization.

Here, we introduce TMEformer, a tumor microenvironment–aware spatial foundation framework. Instead of modeling tumor cells in isolation, TMEformer explicitly incorporates spatial information from high-resolution ST data to jointly learn the interactions of tumor-intrinsic transcriptional programs and surrounding microenvironmental context. For each tumor cell, neighboring cells (conceptually like dynamic spatial niche) are aggregated through attention-based mechanisms to generate a compact context token summarizing the local TME, which is subsequently integrated with the embedding representation of corresponding tumor cell and then processed by a shared transformer encoder. Based on this design, versatileISP strategies and tasks are implemented to discover both tumor-intrinsic and TME-derived dependencies, enabling cellular ecosystem-aware causal inference. Moreover, the learned TME representations yield interpretable embeddings that capture local tissue architecture and support downstream stratification analysis. Together, TMEformer establishes a generalizable framework for modeling tumors as spatially coupled, perturbable ecosystems and provides a foundation for mechanistic investigation of tumor evolution.

## Results

### 1. A TME-aware Spatial Transformer Model for High Resolution Gene Expression Prediction

To model the interplay between tumor cells and their neighboring microenvironment, we introduce a tumor microenvironment-aware single-cell transformer modeling framework and train it on high-resolution spatial transcriptomic data (**Fig. 1a**). Specifically, a TME-CEM (TME Context Encoding Module) is designed to enable integration of each tumor cell’s intrinsic transcriptional profile with signals from its immediate spatial microenvironment. For a given tumor cell, the module first represents the initial embeddings of its spatially neighboring TME cells encoded by overall transcriptome and annotated cell types. These TME embeddings are processed through two sequential attention layers to model intercellular communications and aggregated into a single context token, which is then fused with the tumor cell’s intrinsic embedding using a fixed weight (**Methods**). Thus, our **TME**- aware spatial Trans**former** (TMEformer) encoder infers context-dependent regulatory drivers of tumor heterogeneity, enabling a systems-level view of the tumor ecosystem. It achieves this by integrating spatial architecture to first model the distinct cellular components surrounding each tumor cell and then orchestrate their complex interplay, thereby advancing far beyond conventional single cell gene regulatory network (GRN) inference.

**Fig. 1.**
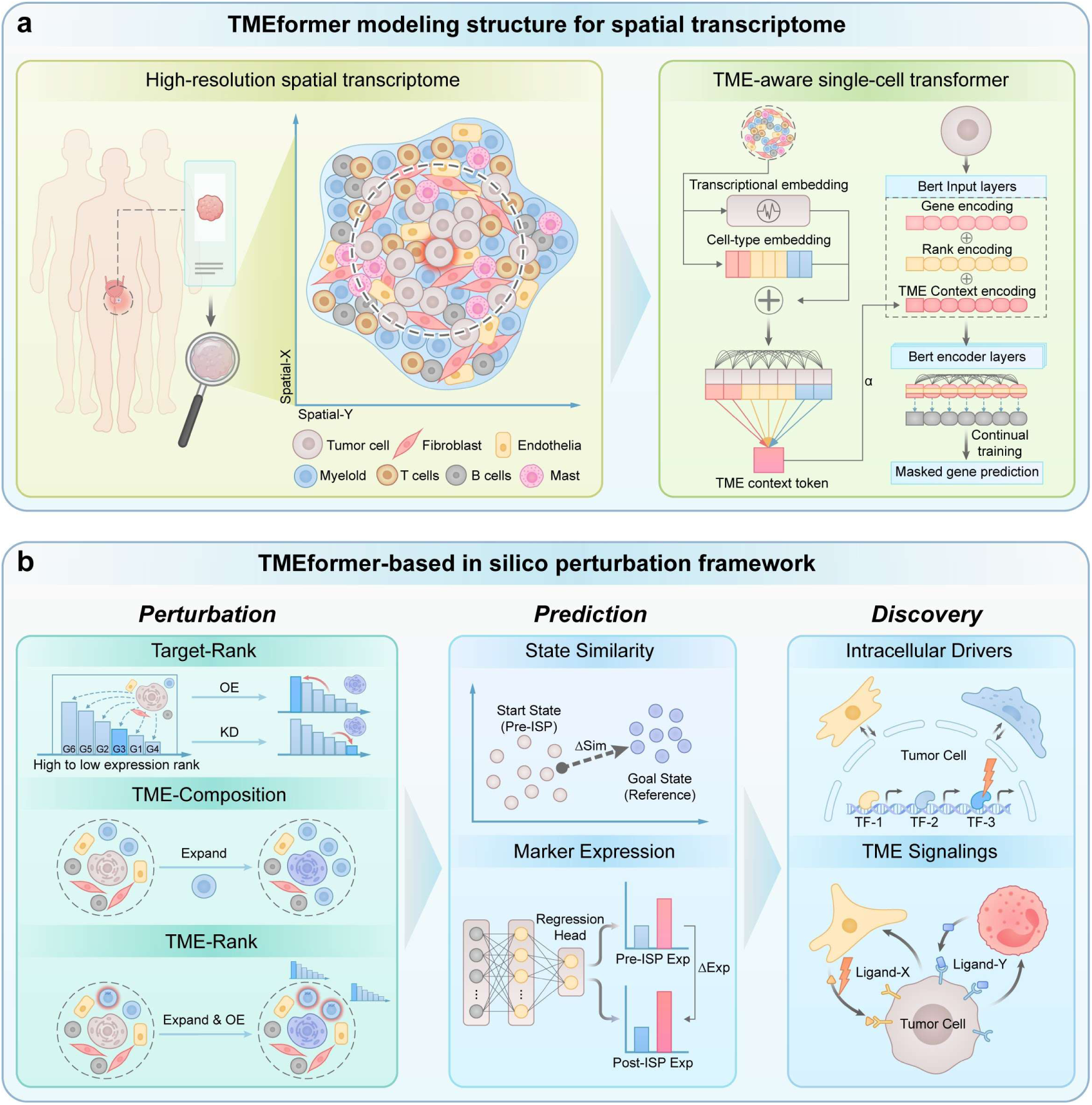
Overview of the TMEformer framework. **a**, The demonstration of TMEformer modeling structure for spatial transcriptome. **b**, TMEformer perturbation framework, comprising perturbation (three ISP strategies), prediction (two predictive tasks), and discovery (intracellular drivers and TME signalings identification).

For TMEformer to achieve optimal performance, high resolution spatial transcriptomic data is essential. For this reason, we acquired clinical spatial transcriptomic data on eight prostate cancer (PCa) specimens spanning key disease stages using the Xenium-5K platform (**Figs. S1a-b**). This platform offers unparalleled subcellular resolution for a panel of 5,000 predefined genes, which we extended by adding customized probes for 51 PCa associated targets (**Supplementary Table 1**). After standard processing and quality control, we obtained spatial profiles for 5051 genes across 1,495,698 cells (**Methods**). Major cell types were annotated using canonical markers, identifying seven main populations: epithelial cells (PCa cells, 49.2%), fibroblasts (24.2%), T cells (9.6%), endothelial cells (7.0%), myeloid cells (6.8%), B cells (2.5%), and mast cells (0.7%) (**Figs. S1c-f**). Subsequently, we performed both tumor-specific and TME-aware continual training on this dataset, enabling the model to adapt to the specific transcriptional landscape of prostate tumors while integrating spatial microenvironmental context. In sample-independent evaluations, TMEformer showed markedly reduced loss compared with the baseline model lacking TME information and ablation analyses across multiple TME-CEM components, including TME inputs and attention-based context encoding, consistently supported the effectiveness of the adopted architecture (**Supplementary Table 2**).

### 2. TMEformer Demonstrates Robust Performance in Predicting Cellular Response under In Silico Perturbation

Cellular responses to signals are shaped by complex microenvironmental interactions that most existing single-cell foundation models fail to capture. By explicitly incorporating spatial context, TMEformer provides a more realistic modelling framework. We next assessed its performance in predicting such responses from different perspectives by various in silico perturbation (ISP) experiments (**Fig. 1b**, **Methods**). Three baseline models were used for benchmarking: the original Geneformer model (GF_PR), its continued-training counterpart on pan-cancer scRNA-seq data (GF_CL), and a version further trained on our PCa spatial transcriptomic data without incorporating spatial TME information (GF_PCa).

Changes in marker gene expression provide an interpretable readout of how ISPs influence cellular phenotypes. To this end, we designed a fine-tuning task to predict gene expression under stimulated perturbations (**Fig. 2a**, **Methods**). Focusing on neuroendocrine prostate cancer (NEPC) – a therapy-resistant state arising from adenocarcinoma through profound molecular reprogramming – we accessed the model’s capability to recapitulate this process, using a panel of canonical NEPC markers (SYP, CHGA, ENO2, and NCAM1) as the molecular readout. After model fine-tuning, we validated its predictions across multiple ISP tasks using established drivers of neuroendocrine differentiation: overexpression (OE) of SOX2^17^ and ASCL1^18^, combined loss of PTEN, RB1, and TP53 (TKO) ^19^, and LIF-LIFR, the top ligand-receptor (L-R) pair for lineage plasticity identified in a recent study^20^. Target gene(s) were shifted to the highest or lowest expression rank to simulate overexpression or knockdown (KD), respectively, with the resulting effect quantified by changes in the endpoint cell embedding (**Methods**). TMEformer showed significant upregulation of NEPC markers following SOX2 or ASCL1 overexpression, as well as TKO perturbation, compared with random perturbations (**Figs. 2b**). Notably, it also exhibits the most robust performance, achieving the best results in three of the four tasks (**Figs. 2c, S2**). Moreover, among all the 15 candidate L–R pairs identified in the previous study^20^, TMEformer successfully predicted over 60% of pairs with significant overall effect on NEPC makers (**Figs. 2d-e**), outperforming the other models evaluated (**Fig. 2f**). Predictive accuracy was better in post-treatment samples (ADT or CRPC) (**Fig. 2g**) relative to treat-naive samples, consistent with the original reported findings that these signalings are particularly relevant under androgen deprivation therapy. More importantly, ligand overexpression in neighboring PCa cells also elicits a significant increase of NE markers in target cells (**Fig. 2h**), further corroborating TMEformer’s ability to integrate microenvironmental signals and propagate their effects to the cell of interest.

**Fig. 2.**
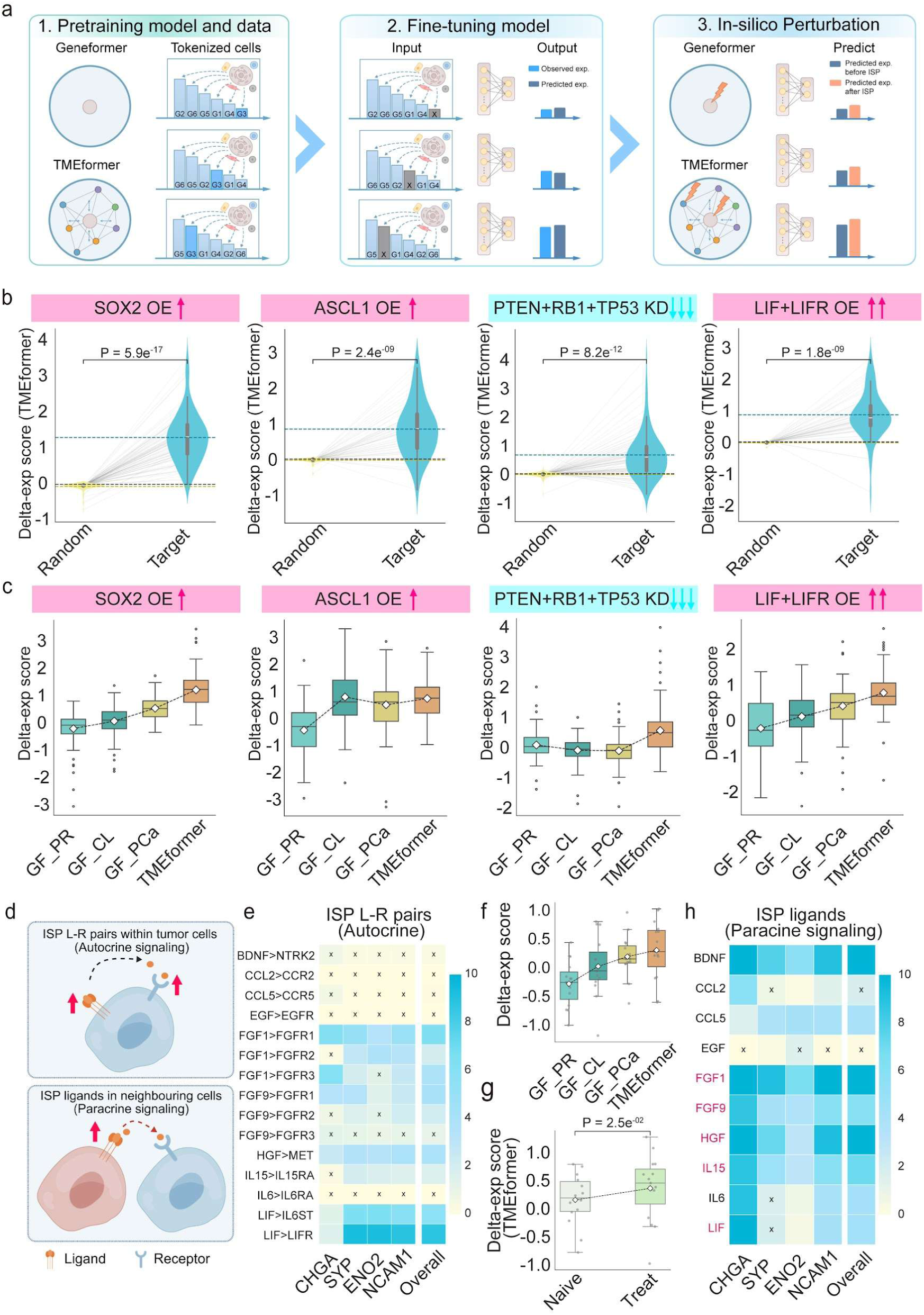
Prediction of ISP effects on NEPC marker expression. **a**, A schematic workflow illustrating the steps of gene expression model fine-tuning and in silico perturbation. Created in part with BioGDP.com. **b**, Violin plots showing the overall delta-exp scores for NEPC markers (CHGA, SYP, ENO2, and NCAM1) under target perturbations (ASCL1 OE, SOX2 OE, PTEN+RB1+TP53 KD, and LIF+LIFR OE) compared with random perturbations, as predicted by TMEformer. Statistical significance was assessed using a one-sided Wilcoxon signed-rank test. Dashed lines indicate group means (blue for Target; yellow for Random) and the zero baseline (black). **c**, Box plots showing the overall delta-exp scores for NEPC markers across models under target perturbations. Diamonds indicate group means. **d**, Schematic illustration of perturbations applied either to ligand–receptor (L–R) pairs within target tumor cells (autocrine signaling) or to ligands in neighbouring tumor cells (paracrine signaling). Created in part with BioGDP.com. **e**, Heatmap showing the effects of 15 L–R pair OE on individual NEPC markers and the overall NEPC marker score, as predicted by TMEformer. Color denotes the −log10(P-value), with “×” indicating non-significant effects (*P* > 0.05). **f**, Box plots showing the overall delta-exp scores for NEPC markers across models under 15 L–R pair OE. Diamonds indicate group means. **g**, Box plots comparing the overall delta-exp scores for NEPC markers under 15 L–R pair OE between treatment-naïve and post-treatment samples predicted by TMEformer. Diamonds indicate group means. Statistical significance was assessed using a Wilcoxon signed-rank test. **h**, Heatmap showing the effects of ligand OE in neighboring PCa cells on individual NEPC markers and the overall NEPC marker score, as predicted by TMEformer. Ligands belonging to significant L–R pairs identified in the Target-Rank ISP analysis (as in **e**) are highlighted in red.

Using the embedding representations of cells at specific disease stages as reference, cellular responses under ISP can be measured by changes in embedding similarity toward that stage. This approach provides a more holistic view of cellular state transitions beyond individual marker genes. Notably, this task can be conducted in a zero-shot manner without model fine-tuning. Castration-resistant prostate cancer (CRPC) represents a critical and therapeutically challenging stage that emerges following androgen deprivation therapy (ADT)^21^. We first benchmarked our models using androgen signaling pathway activation, a hallmark of CRPC^22^. With the CRPC sample from our in-house cohort as the reference state and the ADT samples designated as perturbation targets, TMEformer predicted a significant transition of ADT-treated cells toward CRPC-like representations in contrast to other baseline models (**Figs. S3a**). In particular, the P021 sample with the lowest AR expression showed the most pronounced perturbations effect (**Figs. S3b-c**), further validating the dependency on AR baseline levels. The TMEformer framework enables predictions of how microenvironmental cues influence tumor cells. Focusing on myeloid derived IL- 23, a factor previously implicated in CRPC progression^23^, we simulated overexpression of its subunits (IL23A and IL12B) and assessed the downstream effects on prostate cancer cells. Consistently, both IL12B and IL23A perturbation significantly shift target cells toward CRPC-like states with clear cell-type specificity, especially for IL12B (**Figs. S3d-g**). TMEformer also recapitulated the promotive effect of fibroblast-derived SPP1 on CRPC progression via in silico perturbation (**Figs. S3h-i**), consistent with previous experimental findings ^24^. Incorporating with further subpopulation analysis of fibroblasts(**Fig. S4**), we then found the simulated expansion of the SPP1^high^ subpopulation induced a stronger shift toward the CRPC state compared to their SPP1^low^ counterparts (**Figs. S3j-k**), validating the ability of TMEformer to model microenvironment-driven intercellular influences across different perturbation modalities.

### 3. Screening of Intrinsic Cell Fate Determinants During Tumor Cell Evolution

Having validated TMEformer’s predictive accuracy, we next leveraged its unique capability to perform large-scale in silico perturbation screening under spatially resolved tumor microenvironmental contexts. To systematically identify intrinsic transcriptional regulators that drive prostate cancer cell state transitions during disease progression, we screened key transcription factors (TFs) for their ability to promote three hallmark processes: neuroendocrine differentiation, castration resistance (**Fig. 3a**), and progression toward higher Gleason grades. For these purposes, a list of 495 human TFs (**Supplementary Table 3**; See **Methods**) that overlapped with the Xenium gene panel was investigated. Then, in-silico overexpression for each TF was conducted and corresponding effects on the three processes were respectively evaluated to pinpoint candidate master regulators.

**Fig. 3.**
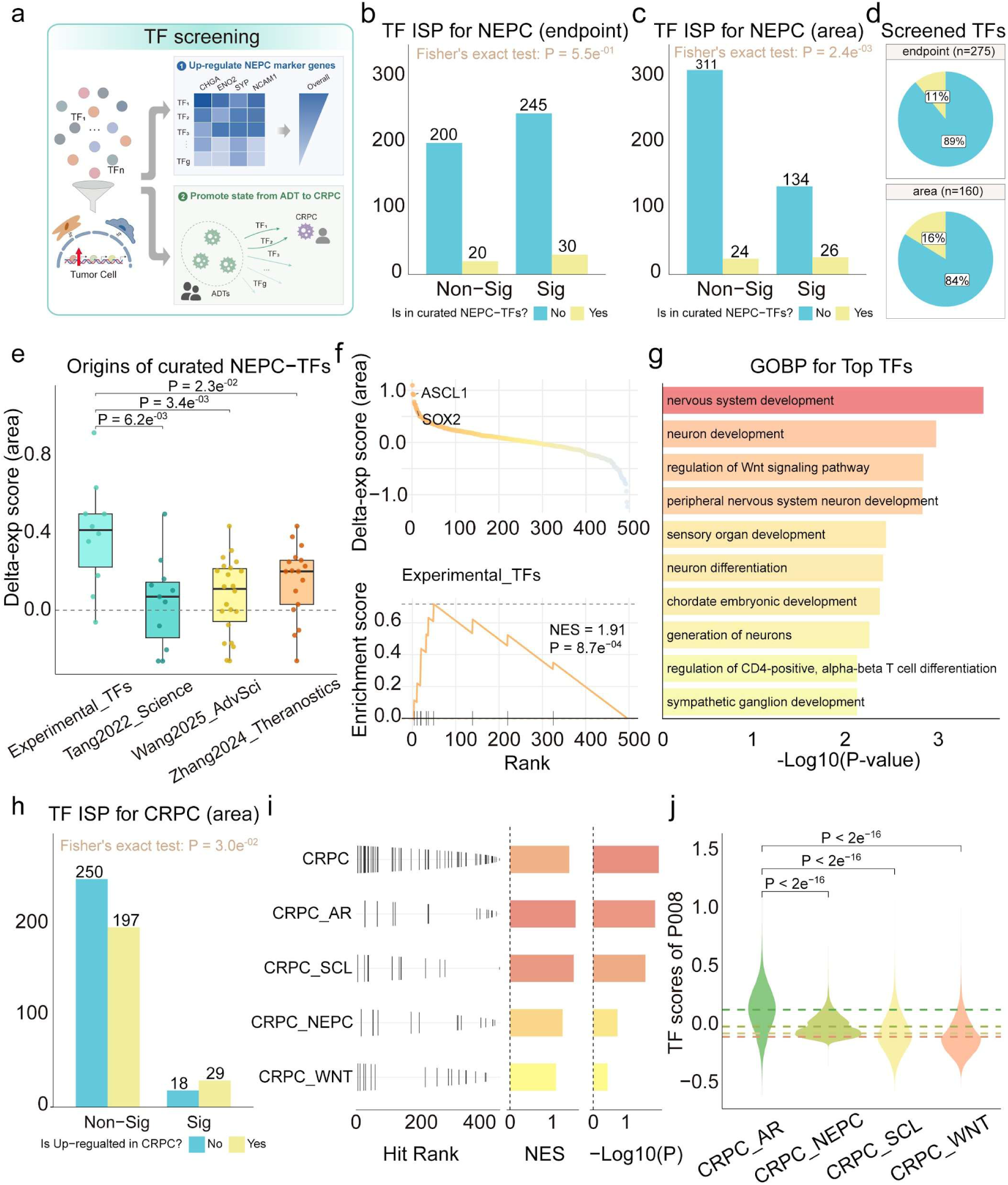
Identification of TFs associated with NEPC marker induction and CRPC state transition. **a**, Schematic overview of TF screening in PCa cells using TMEformer. Candidate TF overexpression perturbations were evaluated based on their ability to upregulate NEPC marker genes or promote cellular state transition from ADT to CRPC. Created in part with BioGDP.com. **b**-**c**, Bar plots showing the numbers of TFs with significant or non-significant predicted effects on NEPC markers. TFs are further stratified by curated NEPC-TFs, and enrichment among significant versus non-significant TFs was assessed using Fisher’s exact test. Perturbation effects were evaluated using endpoint-based (**b**) and area-based (**c**) delta-exp scores. **d**, Pie charts showing the proportions of curated NEPC-TFs among screened TFs identified using endpoint-based and area-based delta-exp scores. **e**, Box plots comparing overall delta-exp area scores for NEPC markers across different TF sets. Pairwise group differences were assessed using Wilcoxon rank-sum tests. **f**, Top: ranked TF plot based on overall delta-exp area scores for NEPC markers. Bottom: enrichment analysis of the NEPC-related TF set (Experiment_TFs). **g**, GOBP enrichment analysis of top-ranked TFs with the highest overall delta-exp area scores for NEPC markers. **h**, Bar plots showing the numbers of TFs with significant or non-significant predicted effects toward the CRPC state. TFs were further stratified by whether they are upregulated in CRPC, and enrichment among significant versus non-significant TFs was assessed using Fisher’s exact test. Perturbation effects were evaluated using delta-sim area scores. **i**, Enrichment analyses of CRPC-related TF sets in TFs ranked by delta-sim area scores toward the CRPC state. **j**, Violin plots showing TF set scores for four CRPC subtype-associated TF sets in PCa cells from the P008 CRPC sample. Statistical significance was assessed using Wilcoxon rank-sum tests. Dashed lines indicate group means.

Given that TFs often function at relatively low abundance levels, the previously used conventional perturbation metrics (endpoint score) may underestimate their effects. We thus developed a scoring strategy based on the area of cumulative perturbation curve (area score) (**Methods**) for more robust evaluation of perturbation effect. We first tested the utility of different scoring metrics on the screening of neuroendocrine differentiation. As expected, while the area-based score yielded results comparable to the endpoint scoring method for individual gene perturbation tasks (**Fig. S5**), it proved particularly advantageous for batch TF screening, as evidenced by its improved accuracy in recognizing curated NE-associated TFs (**Figs. 3b-d, Supplementary Table 4**). Moreover, experimentally validated TFs exhibited stronger perturbation effects and were more strongly enriched at the top of the prediction ranking compared to computationally inferred ones (**Figs. 3e-f, S6**) using the score strategy. Consistently, top ranked TFs were significantly associated with neuron-related and neurodevelopmental pathways (**Fig. 3g**). Similarly, key TFs identified by TMEformer in driving CRPC progression were significantly enriched among genes highly expressed in CRPC relative to other TFs (**Fig. 3h**). In addition, ranked enrichment analysis showed that CRPC-associated TFs were prominently enriched, particularly those related to the AR subtype (**Fig. 3i**), consistent with the elevated expression scores observed for AR-subtype TFs in the CRPC sample within our cohort (**Fig. 3j**).

In parallel, we also screened for drivers that could facilitate PCa progression toward higher Gleason grades^25, 26^. Leveraging the treat-naïve sample P020 with spatially well-defined Gleason 3 (G3) and Gleason 4 (G4) regions (**Fig. S7a**), we identified 63 TFs with significant ISP effect. (**Fig. S7b**). Among them, 12 TFs consistently drove the cellular states toward higher Gleason grades in available treatment-naive samples (**Fig. S7c**). SRF ranked first with the highest perturbation score, and its elevated expression has been strongly linked to prostate cancer progression and castration resistance ^27^. Notably, TWIST1 was involved in multiple enriched pathways, particularly those associated with lipid metabolic processes (**Fig. S7d**), which have been increasingly recognized as a key feature of prostate cancer progression^28, 29^. In addition to the robust perturbation effects (**Figs. S7e-g**), analyses of the TCGA and MSKCC cohorts found that TWIST1 expression was positively correlated with disease status, pathological stage, and Gleason grade (**Figs. S7h-i**), and that elevated TWIST1 expression was associated with prostate cancer recurrence, supporting its clinical relevance (**Figs. S7j-k**). Together, these results suggest that TWIST1 may act as a potential transcriptional driver promoting PCa progression.

### 4. TMEformer Enables Systemic in silico Perturbation of the TME factors

Beyond intrinsic programs, we next leveraged the TMEformer perturbation framework to dissect how extrinsic TME alterations influence prostate cancer cell states by performing systematic ISPs on TME composition and ligands (**Fig. 4a**). Focusing on the NEPC phenotype, compositional perturbations revealed that myeloid cells exerted the most pronounced effect on NE markers compared with other TME cell types (**Figs. 4b-c**). However, two samples showed minimal response and were thus stratified into the low-response (Low-Resp) group, with the remaining samples classified as the high- response (High-Resp) group (**Fig. 4d**). Despite comparable expression levels of canonical myeloid markers such as CD68, CD14 and CSF3R, the High-Resp group displayed higher expression of MRC1 and STAT6, markers associated with immunosuppressive phenotypes, whereas the Low-Resp group was linked to pro-inflammatory signaling, including interferon-γ and TNF pathways (**Figs. S8a-c, 4e**). Further correlation analysis revealed that myeloid cells in the High-Resp group were associated with NE progression, while those in the Low-Resp group showed the opposite trend (**Figs. 4f, S8d**). Correlation-ranked genes revealed significant enrichment of neutrophil-associated signatures, highlighting a potential contribution of N2-like neutrophils to the observed perturbation responses (**Figs. 4g, S5e**).

**Fig. 4.**
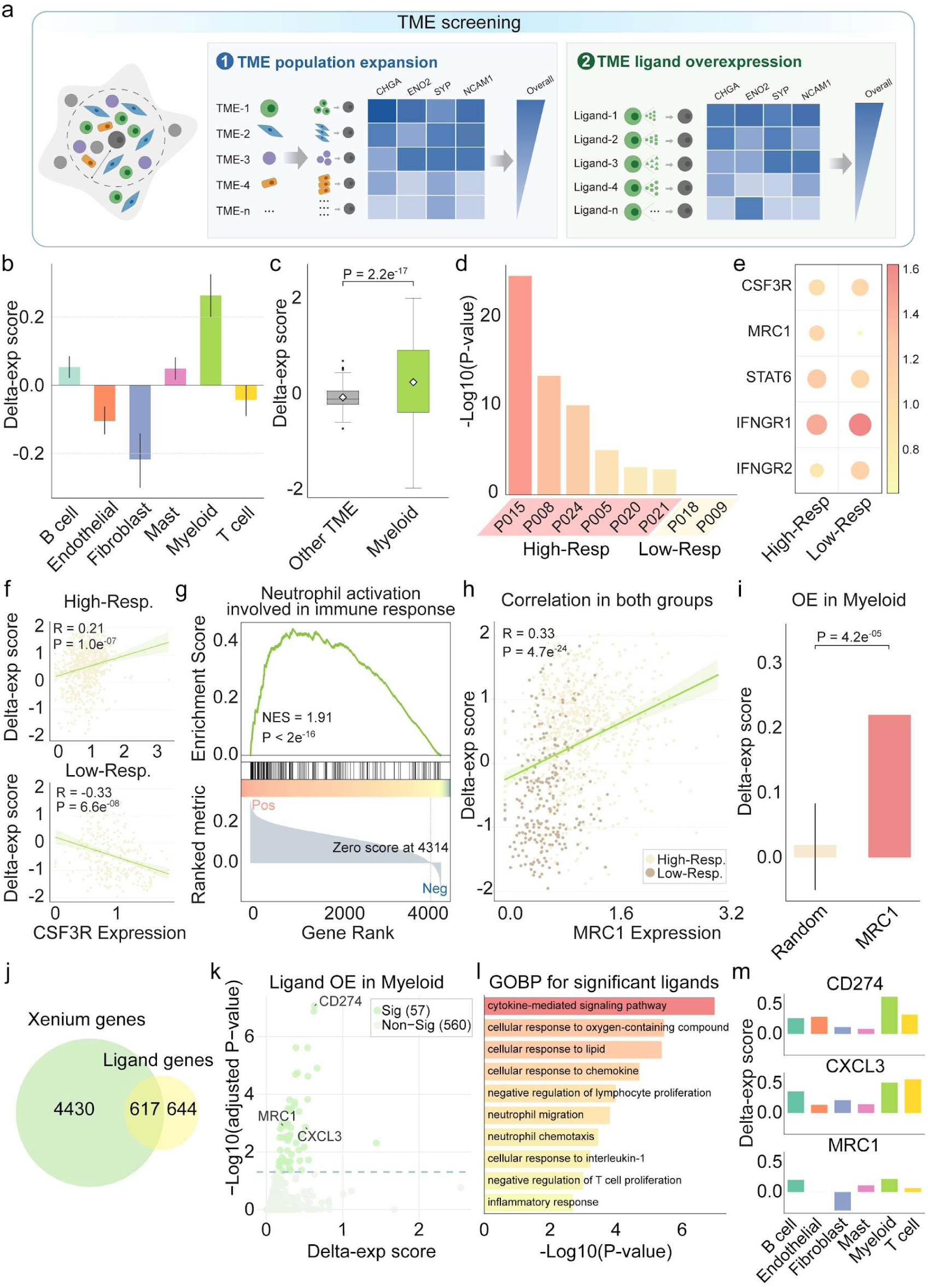
Perturbation in the tumor microenvironment that drives NEPC progress. **a**, Schematic overview of TME factor screening using TMEformer. Expansion of individual TME cell populations was first evaluated for their effects on NEPC marker expression, followed by ligand perturbation screening within pinpointed TME cell types to identify key ligands. **b**, Bar plots showing the overall delta-exp scores for NEPC markers of expanding TME compositions in PCa cells; error bars indicate 95% confidence intervals. **c**, Boxplots comparing the overall delta-exp scores for NEPC markers of expanding myeloid cells and other TME cells. Significance assessed by a one-sided Wilcoxon signed-rank test. **d**, Bar plots show the significance of expanding myeloid cells in each sample. Samples with significant ISP effects on expanding myeloid cells are classified as High-Resp, and the remaining samples as Low-Resp. **e**, Dot plot comparing gene expression in myeloid cells between High-Resp and Low-Resp. Color indicates mean expression, and size indicates the percentage of expressing cells. **f**, Spearman correlation between ISP scores and CSF3R expression in High-Resp (top) and Low-Resp (bottom) groups. **g**, Enrichment analysis of the neutrophil activation–related pathway in genes ranked by correlation with ISP scores in High-Resp group. **h**, Spearman correlation between ISP scores and CSF3R expression with color indicating groups of High-Resp and Low-Resp. **i**, Bar plots showing the ISP scores of MRC1 OE and random OE in myeloid cells on NEPC markers. **j**, Venn diagram showing the overlap between the Xenium genes and collected ligands. **k**, Scatter plots showing the OE effect of ligands in myeloid cells on NEPC markers, with the x-axis indicating the ISP scores and the y-axis indicating significance relative to random perturbations. Ligands with mean ISP scores below 0 are omitted. **l**, GOBP pathway enrichment analysis for ligands with significant ISP effects. **m**, Bar plots showing the mean ISP scores of CD274, CXCL3, MRC1 OE in myeloid cells and other TME components for NEPC markers expression; error bars indicate 95% confidence intervals.

Interestingly, MRC1 expression displayed positive correlation in both groups (**Fig. 4h**) and in-silico overexpression further demonstrated that MRC1 in myeloid cells promotes NE-like features in PCa cells (**Fig. 4i**). Analysis of scRNA-seq from mouse models^19^ revealed an increased proportion of myeloid cells in late-stage NE, accompanied by elevated MRC1 expression, consistent with a progressive enrichment of immunosuppressive myeloid states during NE evolution (**Figs. S9a-c**). To screen more mediators underlying myeloid-driven NE induction, we next performed systematic ligand perturbation (**Supplementary Table 5**; **Methods**) and 57 significant candidates—including MRC1—were identified (**Figs. 4j-k**). Enrichment analysis indicated they were related to cytokine responses, neutrophil migration, and T cell suppression (**Fig. 4l**), which were also observed in myeloid cells from late-NE (**Fig. S9e**). For instance, CD274, a key immunoregulatory molecule in myeloid cells such as tumor-associated macrophages (TAMs) and neutrophils (TANs)^30, 31^, ranked among the ligands with the most significant perturbation effects in a cell type-specific context (**Figs. 4k, 4m**). Its expression was positively correlated with the myeloid expansion-induced NE effect (**Fig. S8f**) and was also upregulated in late-stage NE (**Fig. S9d**). Similarly, CXCL3, a potent neutrophil chemoattractant^32, 33^, exhibited comparable effects (**Figs. 4k, 4m, S8g, S9d**), further supporting a role for neutrophil-associated signaling in mediating NE progression.

In contrast, fibroblast expansion exerted the opposite effect on NE marker expression, with only two samples exhibiting significant responses (**Figs. 4b, S10a**). Differential gene analysis linked the High-Resp group to extracellular matrix (ECM) organization, wound healing, and immunosuppression (**Fig. S10b**). CAF subtype^34^ analysis showed enrichment of immunosuppressive CAF_S1 and multiple myCAF programs in the High-Resp group, whereas normal fibroblast and detox_iCAF signatures were enriched in the Low-Resp group (**Fig. S10c**). Correlation analysis demonstrated that the ECM-associated myCAF signature (ecm_myCAF) was most positively correlated with fibroblast expansion–induced ISP effects, whereas the detox_iCAF signature showed the strongest negative correlation, aligns with the minimal response observed in the Low-Resp group (**Fig. S10d**). Consistent with prior studies^35, 36^ implicating ECM remodeling in NE progression, ligands identified from ISP analyses—including HGF, PDGFC, and COL4A5—were enriched for ECM-related functions (**Figs. S10e–f**). Notably, complement components C4A and C4B were among the top candidates, with expression positively correlated with compositional ISP effects (**Figs. S10g–h**). Moreover, C4 expression was significantly upregulated in fibroblasts from late-stage NE mouse samples (**Fig. S10i**). Prior studies have also linked complement signaling to immune regulation and ECM remodeling within the tumor microenvironment^37, 38^, supporting its potential role in mediating the observed fibroblast-associated effects.

For the TME factors contributing to CRPC progression, in-silico expansion indicated that fibroblasts and T cells exert substantial promotive effects (**Fig. 5a**). Notably, the P009 sample displayed the strongest response to T cell expansion, which may be linked to its higher expression of FOXP3 (**Figs. 5b-c**), a master regulator of regulatory T cells (Tregs). Consistent with this observation, we further identified several Treg-associated ligands (CCL22^39^, BTLA^40^ and EBI3^41^) that exhibited significant ISP effects upon overexpression (**Fig. 5d**). Among these, BTLA is an inhibitory immune checkpoint that can be exploited by tumors to suppress anti-tumor immunity. ^42, 43^. We found its expression was positively correlated with FOXP3 and associated with poorer progression-free interval (PFI) in PCa cohorts (**Figs. 5e-f, S11**), suggesting a potential role of BTLA-mediated Tregs in promoting castration resistance. In contrast, myeloid cells exhibited an opposite effect particularly in P009 and P024 samples (**Figs. 5a-b**), where their deletion increased the state-shifting scores (**Fig. 5g**). Further analysis showed that CXCL9, CXCL10, and CXCL11—canonical markers of M1-like tumor-associated macrophages (TAMs)^44^—were highly expressed in myeloid cells from P009 and P024 but showed lower expression in P021 and the CRPC (P008) sample (**Fig. 5h**). This suggests that M1-like myeloid programs may exert protective effects against CRPC. Significant candidates from ligand screening analyses were enriched in immune-related signatures, among which TGFβ signaling, a hallmark of M2 macrophage polarization implicated in CRPC ^24^, is the most prominent ^45^ (**Fig. 5i**). For instance, TGFB3 with significant perturbation effects specifically in myeloid cells (**Fig. 5j**) has been reported to be closely associated with CRPC ^46^. In addition, four of the top ligands were directly related to M2-like TAMs, including CCL22 and CCL17 (**Figs. 5k-l**), which function as potent chemoattractants for CCR4+ Tregs ^47^. Collectively, these findings support the involvement of the M2-TAM and Treg cells as a potential contributor to CRPC progression (**Fig. 5m**).

**Fig. 5.**
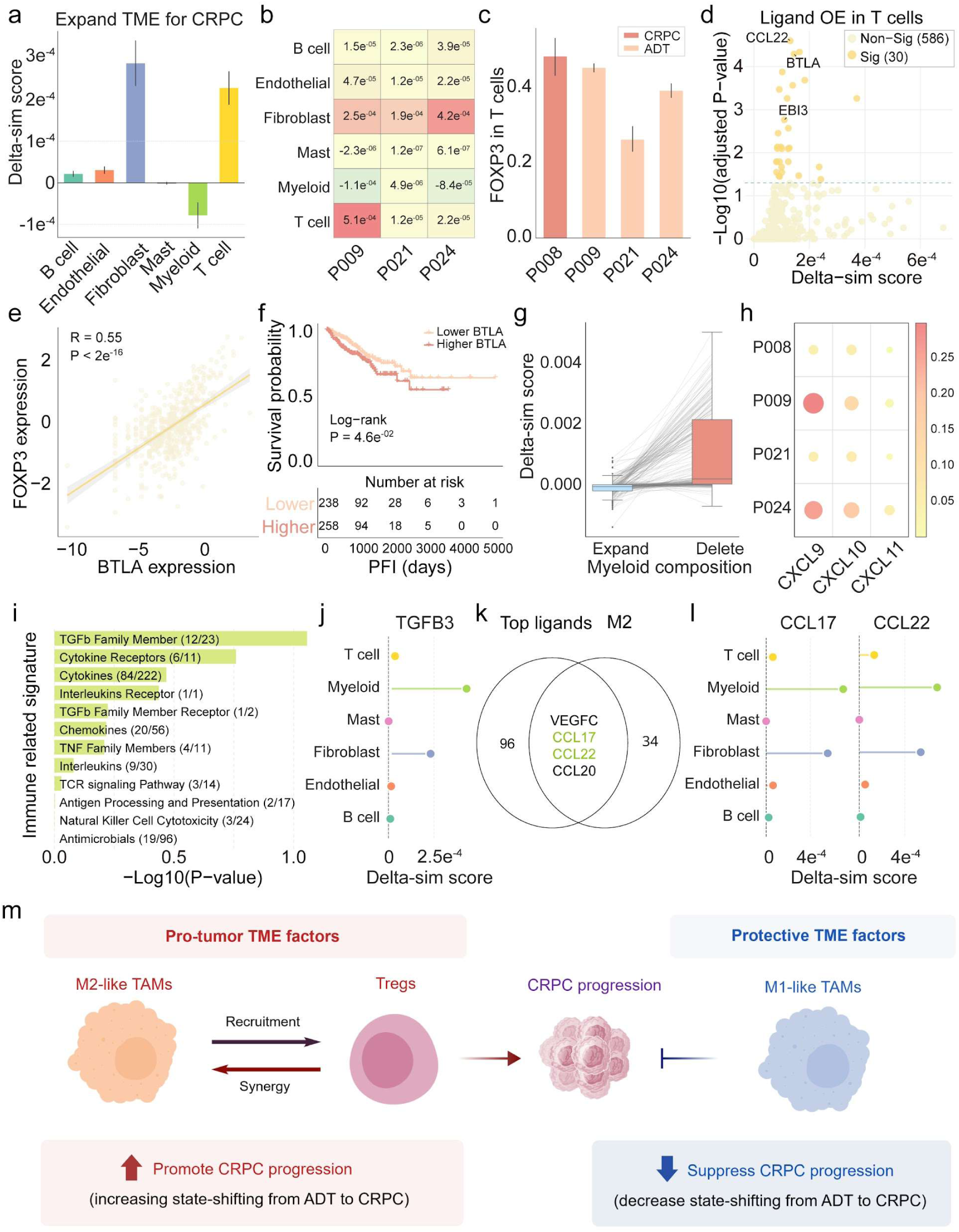
Perturbation in the tumor microenvironment that modulates ADT-to-CRPC transition. **a**, Bar plots showing the delta-sim scores of expanding TME compositions for ADT PCa cellstoward CRPC state; error bars indicate 95% confidence intervals. **b**, Heatmap summarizing the mean delta-sim scores for individual ADT samples toward CRPC state under expansion of different TME cell types. **c**, Bar plot showing FOXP3 expression in T cells across CRPC and ADT samples; error bars indicate 95% confidence intervals. **d**, Scatter plots showing the OE effect of ligands in T cells, with the x-axis indicating mean ISP score toward CRPC state and the y-axis indicating significance relative to random OE. Ligands with mean ISP scores below 0 are omitted. **e**, Spearman correlation between BTLA and FOXP3 expression in TCGA-PRAD cohort. **f**, Kaplan–Meier survival analysis for progression-free interval (PFI) in TCGA-PRAD patients stratified by median BTLA expression. **g**, Box plots showing delta-sim scores toward CRPC state for ADT PCa cells following simulated myeloid expansion or deletion. **h**, Dot plot comparing CXCL9, CXCL10, CXCL11 expression in myeloid cells across CRPC and different ADT samples. Color indicates mean expression, and size indicates the percentage of expressing cells. **i**, Bar plot showing significance between ligands with significant ISP effects and immune-related gene signatures. **j**, Bar plots showing the significance toward CRPC state for ADT PCa cells of TGFB3 OE relative to random genes OE in fibroblasts and other TME components. **k**, Venn diagram showing overlap between the top 100 ligands with highest delta-sim scores and genes associated with M2 phenotypes. **l**, Bar plots showing the mean delta-sim scores toward CRPC state for ADT PCa cells of CCL17 and CCL22 OE in myeloid cells and other TME components; error bars indicate 95% confidence intervals. **m**, Schematic summary of TME factors influencing CRPC progression. Created in part with BioGDP.com.

### 5. TME embedding captures spatial organization and pathological phenotypes

By extracting the learned TME context token embedding (TME-emb) of PCa cells from input layer of TMEformer, we evaluated their utility for tumor stratification and representation from intrinsic transcriptional profiles (EXP-emb) was used as a baseline comparison. Based on the results of unsupervised clustering, we further introduced two metrics—the TME Clustering Index and the Gleason Consistence Index—to quantify spatial coherence and pathological relevance, respectively (**Methods**). As expected, TME-informed clustering consistently showed higher spatial and Gleason coherence across the three treatment-naïve PCa samples with clear Gleason annotation (**Figs. 6a-l**).

**Fig. 6.**
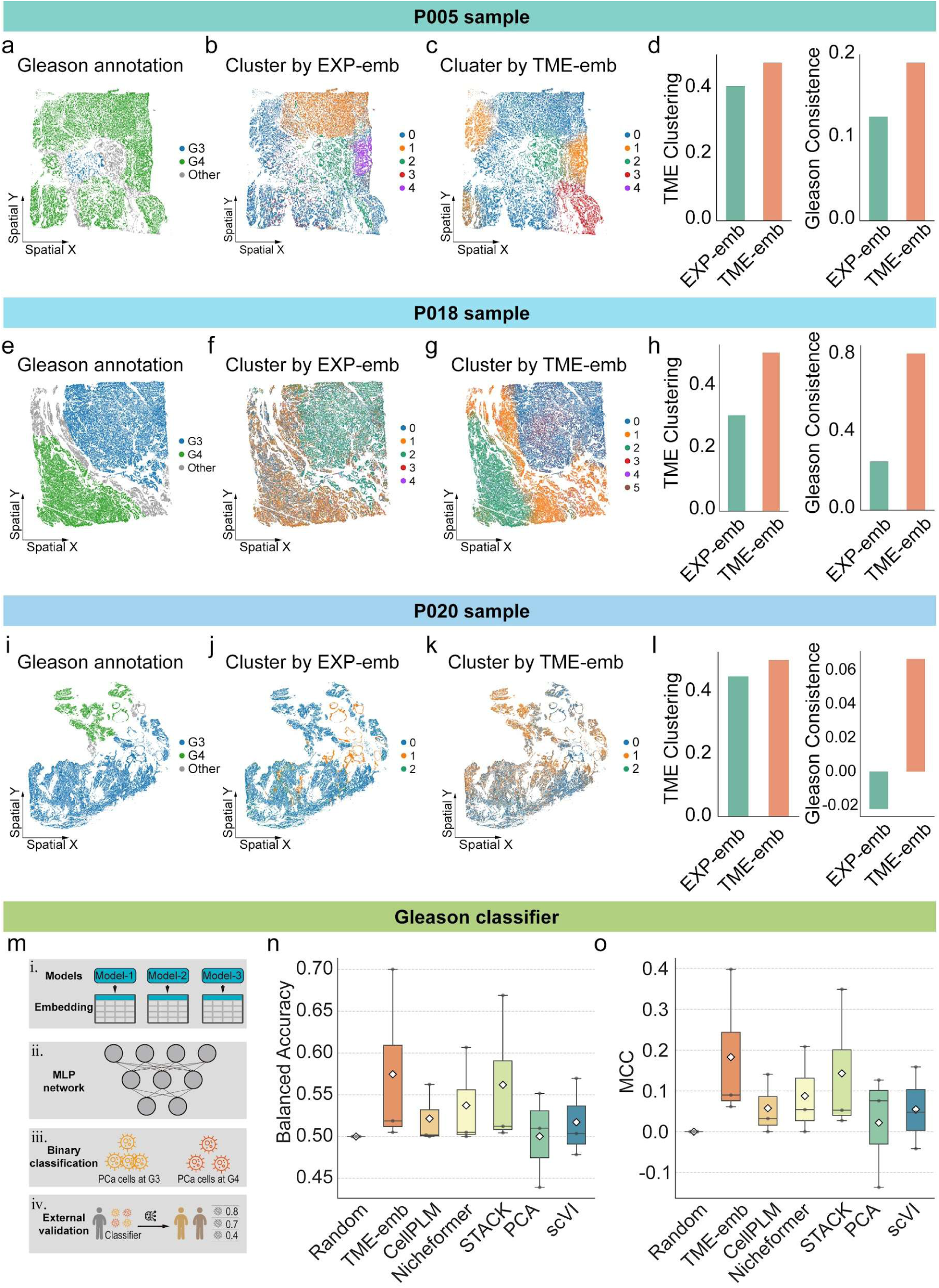
Comparison of expression- and TME-based embeddings for spatial clustering of PCa cells. **a**-**c**, Spatial maps of PCa cells from sample P005, colored by Gleason annotation (**a**), clustered using expression embedding (EXP-emb) (**b**), or clustered using TME embedding (TME-emb) (**c**). **d**, Comparison of the TME Clustering Index and Gleason Consistency Index for EXP-emb– and TME-emb–based clustering in sample P005. **e**-**h**, as in (**a**-**d**), but for sample P018. **i**-**l**, as in (**a**-**d**), but for sample P020. **m**, A schematic workflow illustrating the steps of Gleason classifier model training and validation. **n**, **o**, Boxplots showing the balanced accuracy (**n**) and MCC (**o**) of model cross validation based on different cell embeddings. Each data point represents the performance on the remaining samples for a model trained on a single sample, with diamonds indicating the mean values.

To further validate the phenotypic relevance of TME-emb in a supervised setting, we designed a modeling pipeline to train Gleason classifiers and evaluate cross-sample generalization based on PCa cell representations (**Fig. 6m**). We benchmarked TME-emb against classical dimensionality reduction methods as well as recently proposed single-cell and spatial foundation models. While all models outperformed random prediction, TME-emb consistently achieved the best performance, followed by STACK, Nicheformer, and CellPLM (**Figs. 6n–o**, **S12**). Notably, the recently published STACK model also incorporates cellular context based on scRNA-seq, but does not leverage the advantage of spatial transcriptome. Together, these results demonstrate that explicitly modeling spatially restricted TME context yields cell representations with enhanced predictive power.

### 6. Generalization of TME-aware Modeling to Different Sequencing Platforms and Cancer Types

Finally, we investigated the generalizability of TMEformer across different spatial transcriptomic technologies and cancer types. Tumor stemness represents a critical determinant of therapeutic resistance and metastatic dissemination across diverse cancer types^48^ and was therefore used as a benchmark. To this end, CD44, a well-established marker of tumor stemness ^49, 50^, was used as a molecular readout. We fine-tuned TMEformer to predict CD44 expression in PCa cells and subsequently examined the perturbation effects of classical stemness-associated transcription factors ^51^, namely OSKM (OCT4/POU5F1, SOX2, KLF4, and MYC). In silico upregulation of OSKM consistently led to a more pronounced increase in CD44 expression compared to other genes, both in our in-house dataset and in external PCa spatial transcriptomics datasets from Xenium and MERFISH platforms (**Fig. S13a**). We next extended this analysis to additional cancer types and found TMEformer also maintained robust predictive performance for CD44 response upon OSKM perturbation in both breast and ovarian cancer datasets (**Fig. S13a**), suggesting that the model captures conserved regulatory mechanisms governing tumor stemness across cancers. Importantly, across all evaluated datasets and conditions, TMEformer generally outperformed baseline models (**Fig. S13b**), further highlighting the effectiveness of incorporating TME context into the modeling of gene regulatory responses. Moreover, we simulated the expansion of distinct TME components to assess their impact on tumor cell stemness. Across cancer types and datasets, fibroblast exhibited a more promotive effect relative to other TME cells (**Fig. S13c**). This observation is in line with prior studies ^52, 53^ linking cancer-associated fibroblasts (CAFs) to the maintenance and enhancement of stem-like phenotypes in tumor cells. Collectively, these findings further support the ability of TMEformer to capture TME–tumor interactions across cancer types.

## Methods

### PCa cohort collection and spatial transcriptomic profiling

Tumor tissues from eight patients with prostate cancer (PCa) were collected and subjected to spatial transcriptomic profiling using the 10x Genomics Xenium platform. The cohort comprised four treatment-naïve (TN) tumors, three tumors obtained following androgen-deprivation therapy (ADT), and one castration-resistant prostate cancer (CRPC) sample. For the four TN samples, Gleason pattern–annotated regions were provided based on clinical pathological assessment. All tissue sections were profiled using the Xenium Prime 5K Human Panel. To enhance the detection of pathways associated with prostate cancer, we additionally designed a custom panel consisting of 51 PCa-relevant genes, resulting in a total of 5,051 genes in the standard assay. Raw Xenium output files were processed using the official Xenium analysis pipeline provided by 10x Genomics. Quality control (QC) filtering was applied to remove low-quality cells with fewer than 100 detected genes, as well as genes expressed in fewer than three cells. After QC, high-quality cells were retained for downstream expression normalization and dimensionality reduction using the Scanpy framework. Unsupervised clustering was performed using the Leiden algorithm, and major cell populations were annotated based on the expression of canonical marker genes referring to published literature^54^. For each cell during model training and prediction, genes were ranked based on their expression levels in descending order, and the resulting gene rankings were converted into token sequences following the processing pipelines of Geneformer ^55^.

### TME-aware model designing

We developed a tumor microenvironment (TME)–aware transformer model for spatial transcriptomics, termed TMEformer. The model introduces a TME context embedding module (TME-CEM) at the input layer, enabling the explicit incorporation of spatially informed contextual information from each cell’s surrounding microenvironment. For each PCa cell, neighboring cells were first identified by selecting the top K spatially nearest cells (K = 10,000), from which k = 256 cells were randomly sampled without replacement as candidate TME contributors. For each selected TME cell, two types of embeddings were computed to form its initial representation. First, a transcriptional embedding was obtained using a pretrained single-cell representation model (GF_CL, described below). Second, a learnable cell-type embedding was assigned to each cell based on its annotated cell-type label. These embeddings were combined via element-wise summation. The resulting TME cell representations were processed through a two-stage attention-based aggregation module. The first stage consisted of a self-attention layer that modeled interactions among TME cell tokens and captured higher-order relationships within the local microenvironment. The second stage applied an attention-based pooling mechanism with a learnable query vector to aggregate information into a single TME context token. The TME context embedding was then integrated into the target PCa cell representation using a fixed weighting factor (α = 0.2), and combined with gene token embeddings and rank-based positional embeddings at the input layer. This combined representation served as the final input to the transformer encoder during model training.

### TMEformer training

Two pretrained single-cell transformer models ^56^ were obtained from Hugging Face (https://huggingface.co/ctheodoris/Geneformer). The gf-12L-95M-i4096 model (hereafter referred to as GF_PR) was pretrained on large-scale human single-cell transcriptomic datasets excluding cancer or immortalized cell lines. The gf-12L-95M-i4096_CLcancer model (GF_CL) was subsequently continual-trained on human cancer-related single-cell dataset, which included limited representation of prostate cancer. In this study, annotated PCa cells were used to perform continual pretraining of the GF_CL model using a masked gene prediction objective. This objective enabled the model to capture tumor-intrinsic gene–gene relationships as well as interactions between cancer cell transcriptional programs and their local TME context. Training was performed with a batch size of 6, with gradient accumulation over 4 steps to achieve a larger effective batch size. The model was trained for a single epoch with a maximum learning rate of 0.0001, using the AdamW optimizer with a weight decay of 0.001. A cosine learning rate scheduler with linear warmup over the first 1% of training steps was applied. For benchmarking, a baseline model (GF_PCa) was trained using the identical dataset and architecture initialized from GF_CL, but without incorporating TME context embeddings, while keeping all other configurations unchanged. Further ablation studies were conducted to evaluate the contribution of different components within the TME context module, such as neighbor selection strategies, initial cell embedding calculation, attention-based aggregation layers, and context weighting. Two independent patient samples were used as the validation set, and performance metrics were averaged over three independent training runs to ensure robustness.

### In Silico Perturbation (ISP) implementation

To systematically simulate perturbations at both tumor-intrinsic and microenvironmental levels, we implemented three in silico perturbation (ISP) strategies based on TMEformer.

#### (1) Target-Rank perturbation

This strategy perturbs tumor-intrinsic gene regulation by modifying the expression ranking of one or multiple target genes within one cell, following the scheme introduced in Geneformer. For each cell, genes are ordered by expression level to form an input token sequence. To simulate gene overexpression (OE) or knockdown (KD), target genes are reassigned to higher or lower ranks within the sequence, respectively. The resulting changes in the cell embedding (output of the transformer encoder) are interpreted as the transcriptional impact of the perturbation.

#### (2) TME-Composition perturbation

To capture microenvironmental influences, we perturbed the cellular composition of the local tumor microenvironment. For each tumor cell, the proportion of a specific TME cell type among its neighboring cells was increased or decreased by resampling cells of the target type, while maintaining a fixed neighborhood size (k). The resulting changes in the final cell embedding were used to assess the impact of TME compositional shifts on tumor cells.

#### (3) TME-Rank perturbation

This strategy simulates gene-level perturbations within specific TME cell types. For OE perturbation, a two-step procedure was applied. First, within the selected TME cell type, non-expressing cells were replaced by cells expressing the target gene via resampling, thereby increasing the fraction of expressing cells. Second, for all cells of this cell type, the target gene was reassigned to the highest expression rank. These combined perturbations modulate both the prevalence and transcriptional level within the TME, enabling assessment of how microenvironmental transcriptional alterations affect tumor cell representations.

### ISP evaluation

To quantitatively assess perturbation effects, we developed two complementary scoring schemes: an endpoint score and an area score. Unless otherwise specified, ISP evaluations reported in this study are based on the endpoint score.

#### (1) Endpoint score

The endpoint score measures the perturbation effect by placing the target gene at an extreme position in the rank-ordered gene sequence (*E*_end_). Specifically, for each perturbation, the target gene is repositioned either to the first position, simulating gene overexpression (OE), or to the last position, simulating gene knockdown (KD). The resulting change relative to original cell (*E*_pre_) is defined as the endpoint perturbation effect. This scoring scheme is applicable across all ISP strategies, including Target-Rank, TME-Composition, and TME-Rank perturbations. For TME-composition perturbations, the perturbation axis is defined by the relative fold change in the proportion of a selected TME cell type (e.g., a value of 2 indicates doubling its proportion, whereas 0.5 indicates reducing it by half). Perturbation-induced changes in prediction outputs are computed similarly to the gene-level ISP metrics.

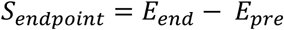

#### (2) Area score

To capture the continuous, gradient-like effects of gene-level perturbations, we additionally introduced an area-based scoring method. Using OE as an example, the target gene is iteratively shifted forward along the rank-ordered sequence in fixed window-size increments (***w***), with each shift representing a single ISP step. At each step, the relative change in gene rank (*r*_0_) is recorded as the x-coordinate, while the corresponding perturbation effect is recorded as the y-coordinate. Notably, for the last perturb (***N***), the target gene is directly repositioned to the endpoint rank if the remaining rank displacement is smaller than predefined window size (***w***). Connecting these points yields a perturbation trajectory curve, and the area under this curve is computed as the area score. A larger area score reflects a stronger cumulative perturbation effect across the entire rank-shifting trajectory, which can reduce stochastic bias introduced by reliance on a single extreme perturbation. Notably, the area score is primarily applicable to Target-Rank perturbations.

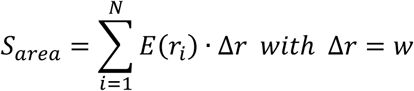

### Predictive tasks for ISP

To explain the biological impact of in silico perturbation on PCa cells, we designed two predictive tasks: state embedding similarity analysis under a zero-shot setting and marker gene expression prediction via model fine-tuning.

#### (1) State embedding similarity

This task evaluates whether a given ISP drives a PCa cell from an initial cellular state (starting state) toward a predefined target state (end state). Under the zero-shot setting, up to 10,000 cells were randomly sampled from the end-state population to compute a mean reference embedding. For each of starting state samples, up to 1,000 cells were randomly selected, and both their original (pre-ISP) and perturbed (post-ISP) embeddings were obtained. Embeddings were extracted from the second-to-last transformer layer for embedding similarity analysis. Cosine similarity was then calculated between the end-state reference embedding and the starting-state embeddings before and after perturbation. The delta similarity score was defined as the difference between post- and pre-perturbation cosine similarity. A positive delta similarity (delta-sim) score indicates that the perturbation shifts the PCa cell representation closer to the end-state phenotype, whereas a negative score suggests divergence from the target state.

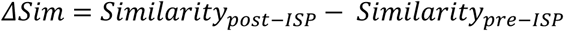

#### (2) Marker expression prediction

This task evaluates the impact of ISP on the expression of one or multiple predefined biological marker genes that characterize specific cellular states. In contrast to the zero-shot state embedding similarity analysis, this task requires model fine-tuning to enable quantitative gene expression prediction. For a given set of marker genes, PCa cells expressing at least one marker were first randomly sampled (up to 100,000 cells). For each selected cell, the marker gene(s) were masked from the expression profile, and the remaining genes were re-tokenized. A multi-output regression head was appended to the model’s output layer to predict the expression values of the masked marker genes. Model optimization was performed by minimizing the sum of mean squared error (MSE) losses across all predicted marker genes. The default hyperparameters were as follows: learning rate = 1 × 10^−4^, weight decay = 0.01, batch size = 16, warmup steps = 200, cosine learning rate scheduler, and 1 epoch. To preserve the pretrained representations and reduce computational cost, the input embedding layers and the top six Transformer encoder layers were frozen during fine-tuning. For single marker prediction, up to 10,000 cells were sampled for evaluation. For multiple markers prediction, up to 5,000 cells for each marker were sampled for evaluation. Model-predicted expression values of marker genes were obtained for each PCa cell before and after perturbation. The delta expression score was defined as the difference between post- and pre-perturbation predicted expression. A positive delta expression (delta-exp) score indicates that the perturbation could increase the expression level of the corresponding marker gene, whereas a negative score implies an inhibitory effect.

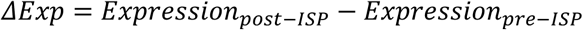

### Background distribution and statistical testing

To assess the statistical significance of in silico perturbation effects, background distributions were generated using randomly selected genes under the same perturbation settings as the target perturbations. For single-gene perturbations, 100 genes were randomly sampled from the available panel genes. Each gene was perturbed in randomly selected cells, yielding 5,000 background perturbations for the embedding similarity task, with 50 cells perturbed per gene, and 10,000 background perturbations for the marker expression task, with 100 cells perturbed per gene. The resulting ISP scores were used to define the background distribution. For multi-gene perturbations involving *m* genes, background distributions were generated by randomly sampling gene combinations of size *m* and applying the same perturbation settings.

To compare target perturbations with the background, each tissue section was divided into square patches of fixed size. Within each patch, the ISP score of each target perturbation was compared with the corresponding background score, defined as the mean ISP score of random gene perturbations in that patch. A one-sided Wilcoxon signed-rank test was then used to assess whether target perturbations produced significantly greater effects than random perturbations. By default, 2,000-µm square patches were used for all-sample ISP analyses, whereas 500-µm square patches were used for few-sample ISP analyses. For external datasets generally containing only one sample, 1,000-µm square patches were used. Statistical significance was defined as *P* < 0.05 by default. For TF and ligand screening analyses, Benjamini–Hochberg (BH) adjusted *P* < 0.05 was used to determine significance.

For marker expression perturbation analyses, overall delta-exp scores were calculated by aggregating effects across marker genes. For individual perturbation analyses, the overall delta-exp score was computed in two steps. First, within each patch, delta-exp scores from target and random perturbations were Z-score normalized separately for each marker gene. Second, the normalized delta-exp scores were averaged across marker genes to obtain an overall delta-exp score for each perturbation. The background overall delta-exp score was defined as the mean overall delta-exp score across random gene perturbations. For batch perturbation analyses, ISP scores from all screened target genes and background perturbations were pooled within the same patch. Scores were then Z-score normalized within each patch separately for each marker gene. For each target perturbation, normalized scores were averaged across marker genes to obtain an overall delta-exp score.

### Transcription factor and ligand gene collection

The gene list of 1,639 human transcription factors (TFs) was obtained from one reliable study^57^, of which 495 genes overlapped with the Xenium gene panel. Four sets of NEPC-related TFs were curated from previous research and 50 of them overlapped with the panel (termed curated NEPC-TFs). One set was curated through a systematic literature review of PubMed with consolidated experimental validation (termed Experimental_TFs). The remaining three sets were inferred from computational analyses based on omics profiling data (termed Tang2022_Science, Wang2025_AdvSci, Zhang2024_Theranostics, respectively). Genes overlapping between the computationally inferred sets and the Experimental_TFs set were excluded. A total of 1,261 ligand genes were integrated from five publicly available databases (CellPhoneDB^58^, CellChat^59^, NicheNet^60^, CellTalk^61^, LRDB^62^), among which 618 genes overlapped with the Xenium panel.

### Enrichment analysis

Pathway enrichment analysis of screened TFs or ligand genes was performed using the python package gseapy with the *enrichr()* function. Pathway gene sets were obtained from Gene Ontology Biological Process (GOBP) database. The background gene set was defined as all candidate TFs or ligands for screening. Rank-based enrichment analysis was conducted using the *fgsea()* function from the fgsea R package. All candidate TFs were ranked according to perturbation scores, and the scoreType parameter was set to “pos” to assess enrichment of phenotype-associated TFs toward the top of the ranked list. Enrichment significance was evaluated based on the normalized enrichment score (NES) and corresponding p-values.

### Public spatial transcriptome datasets of PCa

Publicly available Xenium-5K spatial transcriptomic datasets were obtained from the 10x Genomics data portal (https://www.10xgenomics.com/datasets), including prostate cancer (PCa), breast cancer (BCa), ovarian cancer (OCa) samples. All public datasets were processed using the same preprocessing, quality control, and cell annotation pipeline as applied to our in-house Xenium datasets. For the PCa Xenium dataset, a total of 110,847 cells were retained after quality control. Major cell types were annotated based on canonical marker gene expression, and PCa epithelial cells, accounting for 67.3% of all cells, were used for marker expression perturbation analysis. For the BCa and OCa Xenium datasets, 175,835 and 1,120,560 cells passed quality control, respectively. After cell-type annotation, epithelial cells (66.6% for BCa and 52.4% for OCa), together with their surrounding tumor microenvironment (TME), were used for perturbation analyses. In addition, a spatial transcriptomic dataset comprising two prostate cancer samples generated using the MERFISH platform (500-gene panel; Vizgen MERFISH FFPE Human Immuno-oncology Data Set, May 2022) was included for cross-platform validation.

### Public scRNA-seq datasets of PCa

The Gao dataset^19^ represented a mouse TPPRC-based NEPC model. Tumor tissues were collected after tamoxifen induction at six time points (W2, M1, M2.5, M3.5, M4.5, and M6), together with wild-type (WT) controls. In this study, the first three time points (W2, M1, and M2.5) were defined as the Early_NE stage, whereas the latter three time points (M3.5, M4.5, and M6) were defined as the Late_NE stage. After quality control, 126,838 cells were retained and annotated into the same major cell types. Gene expression scores for predefined gene lists within specific cell populations were calculated using the *scanpy.tl.score_genes()* function. Differential gene expression analysis between stages across different cell types was performed using *scanpy.tl.rank_genes_groups()* with the Wilcoxon rank-sum test. Genes with log2 fold change (log2FC) > 0 and adjusted P-value (padj) < 0.01 were defined as highly expressed genes.

### Public bulk RNA-seq of PCa

RNA-seq expression profiles of the TCGA-PRAD cohort (tcga_RSEM_gene_tpm.gz), together with corresponding clinical (including Gleason score, T/N stage) and survival information, were downloaded from the UCSC Xena database^63^. Kaplan–Meier survival analysis and visualization were performed using the survival and survminer R packages, with patients dichotomized by either the median value or an optimal cutoff. In addition, gene expression data as well as clinical information from the MSKCC cohort^64^ were used for independent validation.

### TME embedding-based clustering

Based on the TME context encoding module of the TMEformer model, a TME context token embedding was computed for each PCa cell. These embeddings (termed TME- emb) were used to perform unsupervised clustering, and the resulting TME-driven clusters were compared with clusters derived solely from the intrinsic transcriptomic profiles (termed EXP-emb) of PCa cells. The comparison focused on spatial coherence and pathological consistency.

#### (1) TME Clustering Index

To evaluate the spatial coherence of clusters, we quantified how spatially proximal cells within the same cluster were, compared with cells from different clusters. Two metrics were used: (i) Silhouette coefficient: Measures the separation between each cell’s assigned cluster and its nearest neighboring cluster. Higher values indicate more distinct and well-separated clusters in spatial space. (ii) iLISI_graph: Assesses the degree of cluster mixing by quantifying the presence of neighboring cells with different cluster labels. Lower mixing (i.e., lower iLISI_graph values) indicates higher spatial purity of clusters. Together, these two metrics form the TME clustering index (Silhouette - iLISI_graph), which reflects how well the clustering captures spatial structure.

#### (2) Gleason Consistency Index

To assess how well the TME-driven clusters correspond to pathological Gleason regions, we computed the Gleason consistency index based on two metrics: (i) Adjusted Rand Index (ARI): Measures the agreement between TME-based clusters and Gleason-labeled regions, adjusted for chance. Higher ARI indicates stronger alignment. (ii) Normalized Mutual Information (NMI): Quantifies shared information between clustering and pathology labels. Higher NMI reflects greater consistency in overall partition structure. Gleason Consistence Index from these metrics (ARI + NMI) jointly evaluate the degree to which TME-derived clustering captures clinically relevant pathological heterogeneity. The above original indexes (Silhouette, iLISI_graph, ARI, NMI) were computed using the scib python package^65^.

### TME embedding-based Gleason classification

A unified Gleason classification pipeline was designed to evaluate the predictive capacity of the learned embedding from different models. In addition to TME-emb, other foundation model embeddings were extracted in a zero-shot manner. scVI was trained with n_latent = 512, n_layers = 2, and dropout_rate = 0.1. PCA was applied after standardization to reduce expression profiles to 512 dimensions using sklearn toolkit. Nicheformer and CellPLM embeddings were extracted using the corresponding default pipelines based on the latest model checkpoints. STACK embeddings were generated using the pretrained bc_large.ckpt model and reduced from 1600 to 512 dimensions via PCA (97.7% variance retained). Random features were sampled from a standard normal distribution. All classifiers shared the same multilayer perceptron architecture (hidden layers: 512–256–128, ReLU, dropout = 0.3). The output layer had two units corresponding to binary Gleason region labels, optimized using the cross-entropy loss. The classifier was trained using the Adam optimizer (learning rate = 0.001, batch size = 128) for three epochs in a cross-sample evaluation setting, where one sample was used for training and the remaining samples were used for evaluation. Performance was assessed using balanced accuracy, Matthews correlation coefficient (MCC), area under the ROC curve (AUC), and precision–recall AUC (PR AUC). For balanced accuracy and MCC, a fixed classification threshold of 0.5 was applied.

### Data and code availability

All public datasets used in this study are publicly available. Xenium spatial transcriptomics datasets were obtained from the 10x Genomics data portal at https://www.10xgenomics.com/datasets, including prostate cancer, breast cancer, ovarian cancer. The MERFISH dataset (Vizgen MERFISH FFPE Human Immuno-oncology Data Set, May 2022) is available from Vizgen. The public single-cell RNA-seq dataset was downloaded from https://ngdc.cncb.ac.cn/omix/release/OMIX001928. TCGA-PRAD data were downloaded from the UCSC Xena database (https://xenabrowser.net/). The tokenized spatial transcriptome datasets generated and analyzed during this study, along with the trained TMEformer model files, are available in the Open Science Framework (OSF) repository at https://osf.io/p3qgu/overview?view_only=093ce25176974361a233252d89896ebe.

The TMEformer source code is publicly available on GitHub at https://github.com/lishensuo/tmeformer. TMEformer is available as a web-accessible platform at http://www.pradcellatlas.com/ for online prediction, analysis, and visualization of virtual perturbation effects.

## Discussion

In this study, we present TMEformer, a tumor microenvironment-aware transformer framework that models tumor cells as components of spatially organized cellular ecosystems. By integrating tumor-intrinsic programs with microenvironmental signals from clinical specimens which capture a rich set of endogenous perturbations such as disease states and tissue niches, TMEformer enables spatially informed in silico perturbation. This approach addresses a central challenge in tumor biology of understanding how perturbations propagate through spatially structured networks to shape cellular states.

Across diverse perturbation settings, TMEformer outperformed Geneformer-derived baselines, particularly in contexts where tumor evolution is tightly coupled to microenvironmental regulation. Notably, the model captures both tumor-intrinsic and non-cell-autonomous effects, allowing perturbations originating from neighboring cells (e.g. ligand signaling or compositional changes) to propagate and reshape tumor cell states. This provides the approach to dissect multi-level regulatory dependencies underlying neuroendocrine differentiation, castration resistance, and tumor progression. Leveraging this framework, TMEformer recovers established drivers and prioritizes clinically relevant candidates, including TWIST1, whose association with disease progression. The model also uncovers non-obvious intercellular dependencies, such as immunosuppressive myeloid states and Treg-associated signaling, demonstrating its ability to reveal mechanisms that are difficult to capture through descriptive spatial analyses. Beyond perturbation prediction, TME-derived embeddings enhance the representation of tumor cell states by embedding them within their local ecological context. This leads to improved spatial coherence and stronger alignment with pathological architecture, indicating that spatial context provides biologically meaningful information not captured by transcriptome-only models. Notably, the generalization of TMEformer across spatial platforms and cancer types suggests that it captures conserved regulatory principles governing tumor–TME interactions.

Despite these advances, several limitations remain. The reliance on predefined spatial neighborhoods and fixed weighting may limit the modeling of long-range or dynamic interactions. In addition, while in silico perturbation enables scalable hypothesis generation, experimental validation is required for newly identified regulators. The current framework is also limited to transcriptomic features and does not yet integrate other spatial modalities such as proteomics or chromatin accessibility. Future work may incorporate adaptive neighborhood modeling, multimodal spatial data integration, and temporally informed frameworks to better capture tumor evolution dynamics. Together, our findings support a shift from cell-centric modeling toward ecosystem-level representations of disease, where cellular behavior emerges from spatially structured interactions. By enabling predictive and perturbation-driven modeling of tumor ecosystems, TMEformer provides a foundation for mechanistic discovery and therapeutic prioritization in cancer.

## Acknowledgements

This study was supported by National Natural Science Foundation of China (92474108, 82373106), Noncommunicable Chronic Diseases-National Science and Technology Major Project (NO.2023ZD0505600, NO.2023ZD0505601), National Natural Science Foundation of China (32200532), Fundamental Research Funds for the Central Universities (YJ202311), and Fundamental Research Funds for the Central Universities (YJ202523).

## Author contributions

S.L., X.W., L.Y., G.Z. and S.C. conceived the study, designed the TMEformer model, performed the analysis, developed the web platform, and drafted the manuscript and figures. P.L. collected clinical samples and contributed to data acquisition. Z.Z., D.L., G.X. and M.Z. contributed to model design and web platform development. Q.C., Y.T. and J.L. participated in data processing and analysis. L.H., B.C., S.O. and J.J. contributed to data interpretation and visualization. S.L., X.W., L.Y., G.Z. and S.C. interpreted the results, revised the manuscript and figures. X.W., L.Y., G.Z. and S.C. jointly supervised the project. All authors reviewed and approved the final version of the manuscript.

