## Supplementary Information for "Mapping Tumor-Microenvironment dependencies with TMEformer: A spatial foundation framework enabling in silico perturbation"

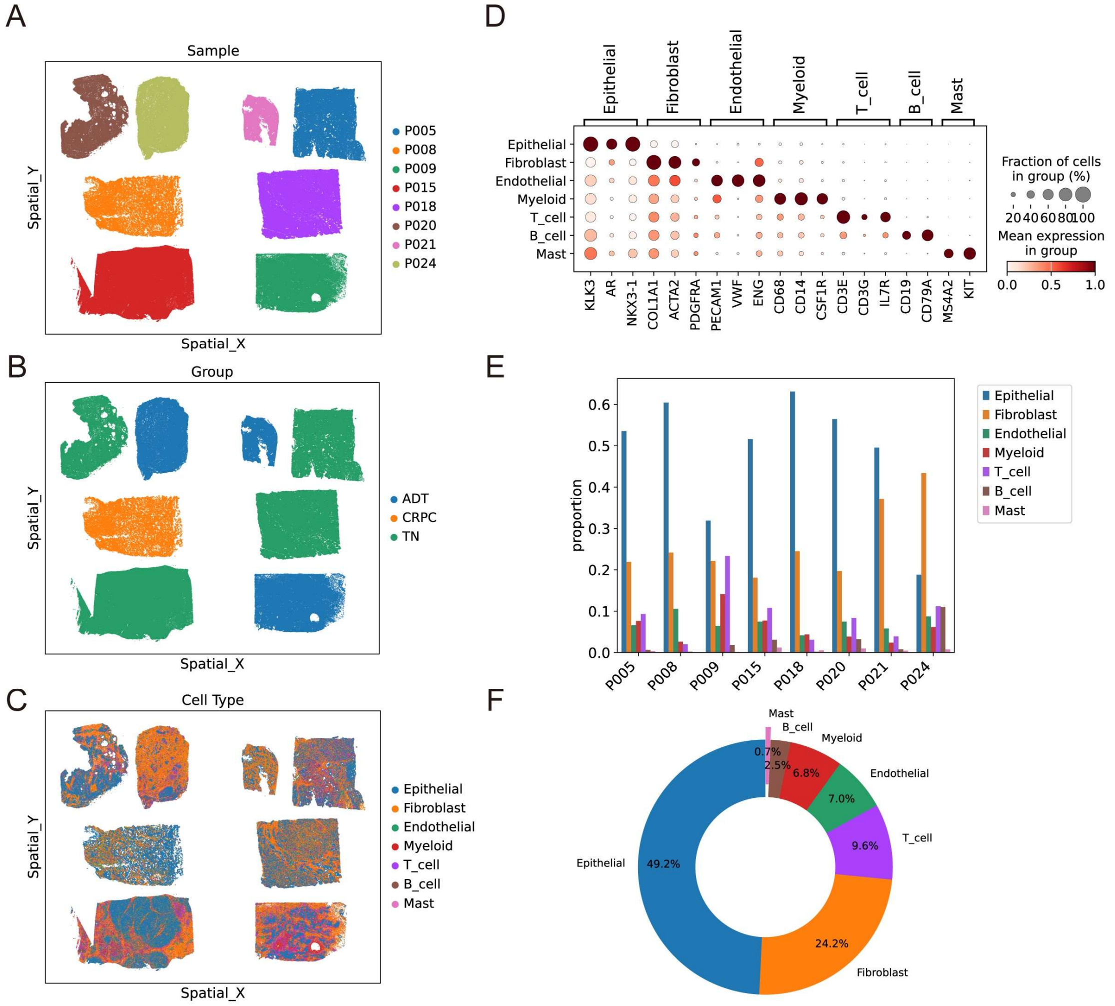


**Supplementary Fig. 1 | Overview of prostate cancer Xenium spatial transcriptomic datasets and cell-type annotation.** (**A**) Spatial distribution of all cells profiled by Xenium in eight PCa tissue samples. Each point represents a single cell, colored by sample identity. (**B**) Same spatial coordinates as in (**A**), with cells colored according to pathological grouping of samples. (**C**) Same spatial coordinates as in (**A**), with cells colored according to annotated cell types. (**D**) Dot plot showing the expression of canonical marker genes used for cell-type annotation. Dot size indicates the fraction of cells expressing each gene, and color intensity represents the average normalized expression level. (**E**) Bar plots summarizing the proportion of each annotated cell type across individual PCa samples. (**F**) Pie chart depicting the overall cell-type composition aggregated across all samples.

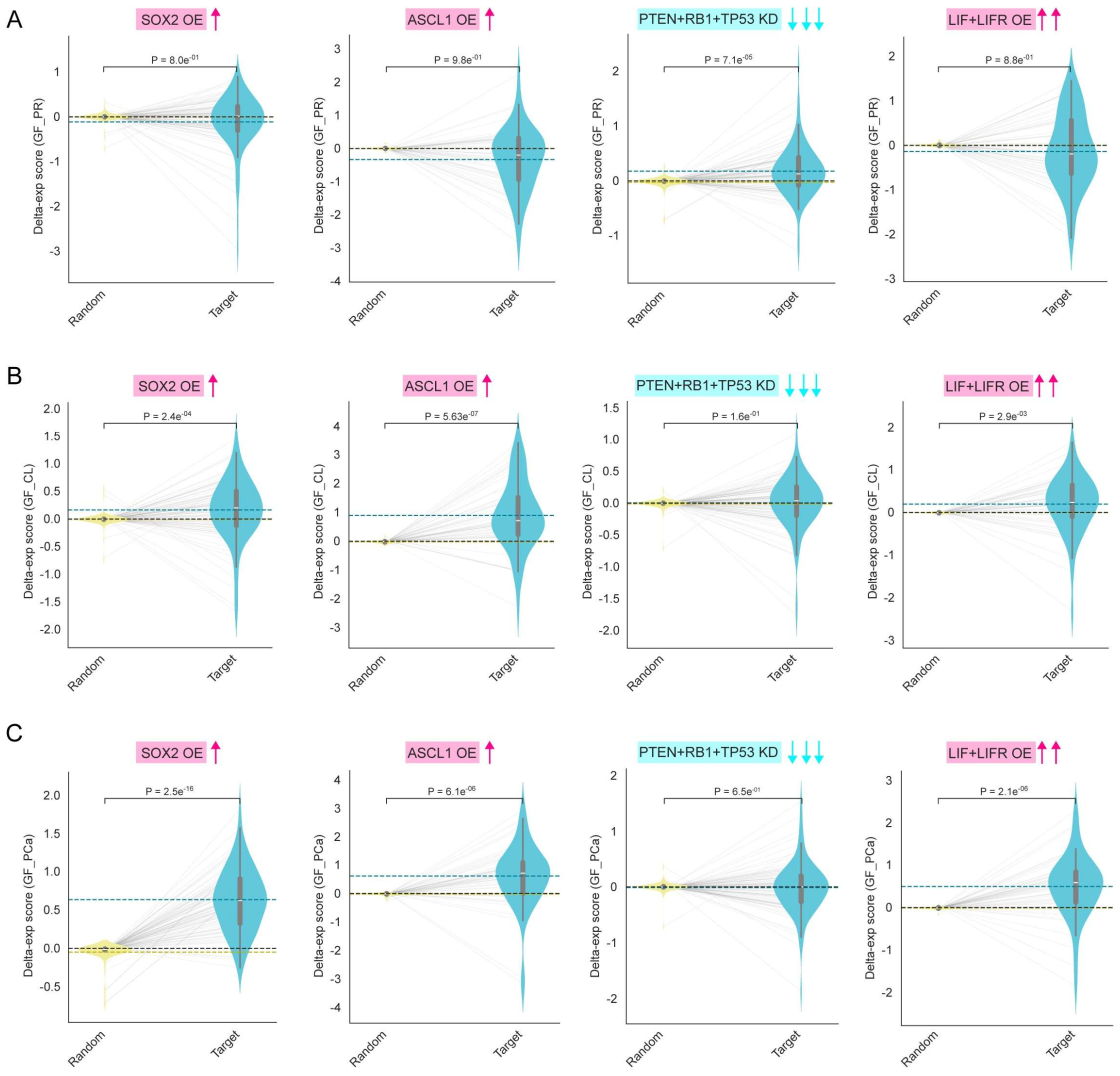


**Supplementary Fig. 2 | External perturbation analyses for NEPC marker expression.** (**A-C**), Violin plots showing the overall delta-exp scores for NEPC markers (CHGA, SYP, ENO2, and NCAM1) under target perturbations (SOX2 OE, ASCL1 OE, PTEN+RB1+TP53 KD, and LIF+LIFR OE) compared with random perturbations, as predicted by the GF_PR (**A**), GF_CL (**B**), and GF_PCa (**C**) models. Statistical significance was assessed using a one-sided Wilcoxon rank-sum test. Dashed lines indicate group means (blue for Target; yellow for Random) and the zero baseline (black).

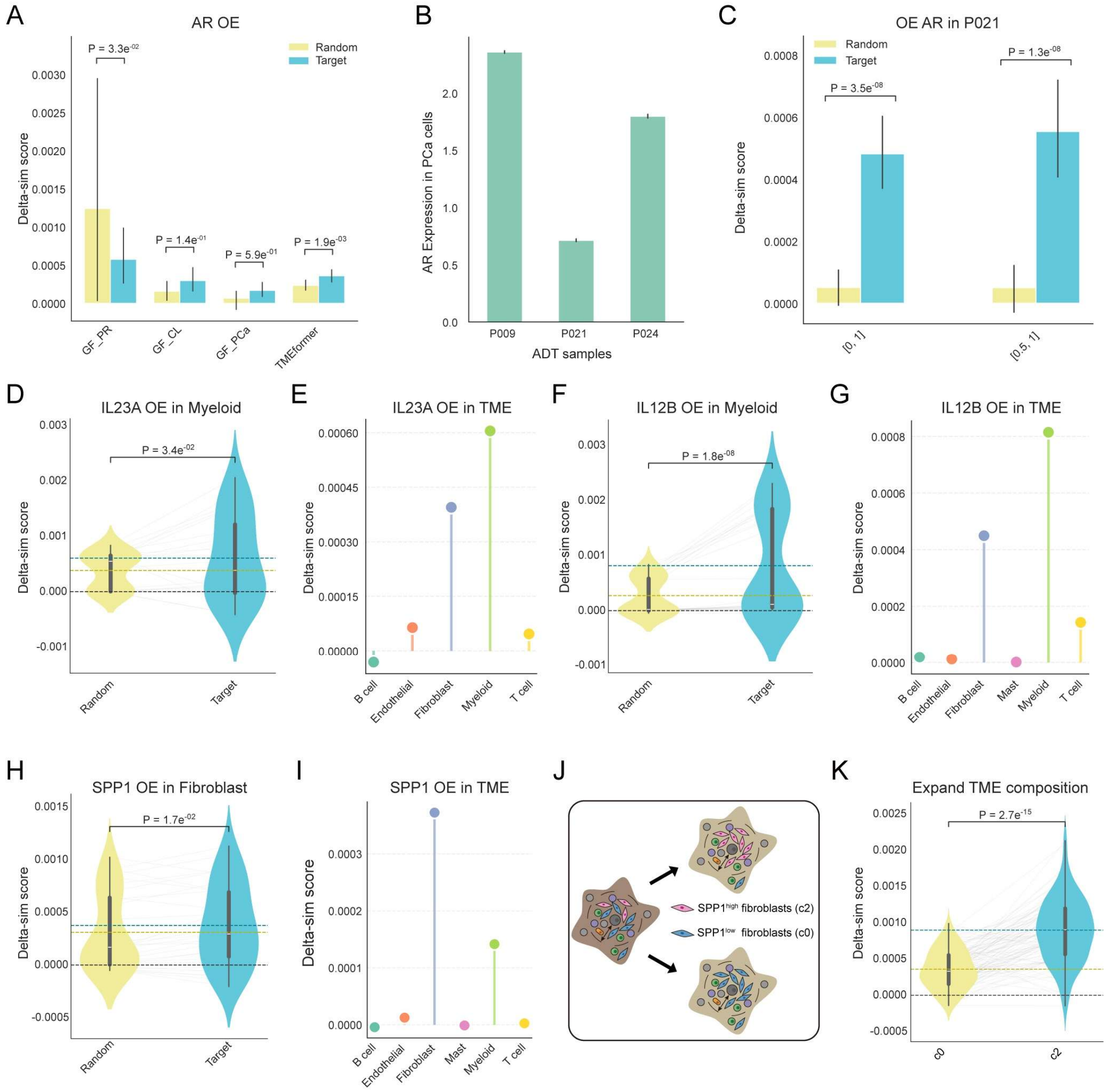


**Supplementary Fig. 3 | Prediction of ISP effects on cellular states towards CRPC state.** (**A**) Bar plots showing the mean delta-sim scores toward the CRPC state for AR OE compared with random perturbations in ADT samples across models; error bars indicate 95% confidence intervals. Statistical significance was assessed using a one-sided Wilcoxon rank-sum test. (**B**) Bar plots showing the mean AR expression levels in PCa cells from different ADT samples; error bars indicate 95% confidence intervals. (**C**) Bar plots showing the mean delta-sim scores of AR OE relative to random perturbations in the P021 sample; error bars indicate 95% confidence intervals. All PCa cells expressing AR are defined as [0,1], whereas low-AR cells ([0.5,1]) correspond to cells in which AR ranks in the bottom 50% of expressed genes. (**D**) Violin plots showing delta-sim scores toward the CRPC state for IL23A OE compared with random perturbations in myeloid cells from ADT samples. (**E**) Dot plots showing the mean delta-sim scores for IL23A OE across different TME cell types. (**F**, **G**), as in (**D**, **E**), but for IL12B OE. (**H**, **I**), as in (**D**, **E**), but for SPP1 OE. (**J**) Schematic illustration of compositional perturbation of different fibroblasts subpopulations. (**K**) Violin plots showing the delta-sim scores toward CRPC state of c2 subcluster expansion (five-fold) relative to c0 subcluster expansion (two-fold) within fibroblasts of P009 sample.

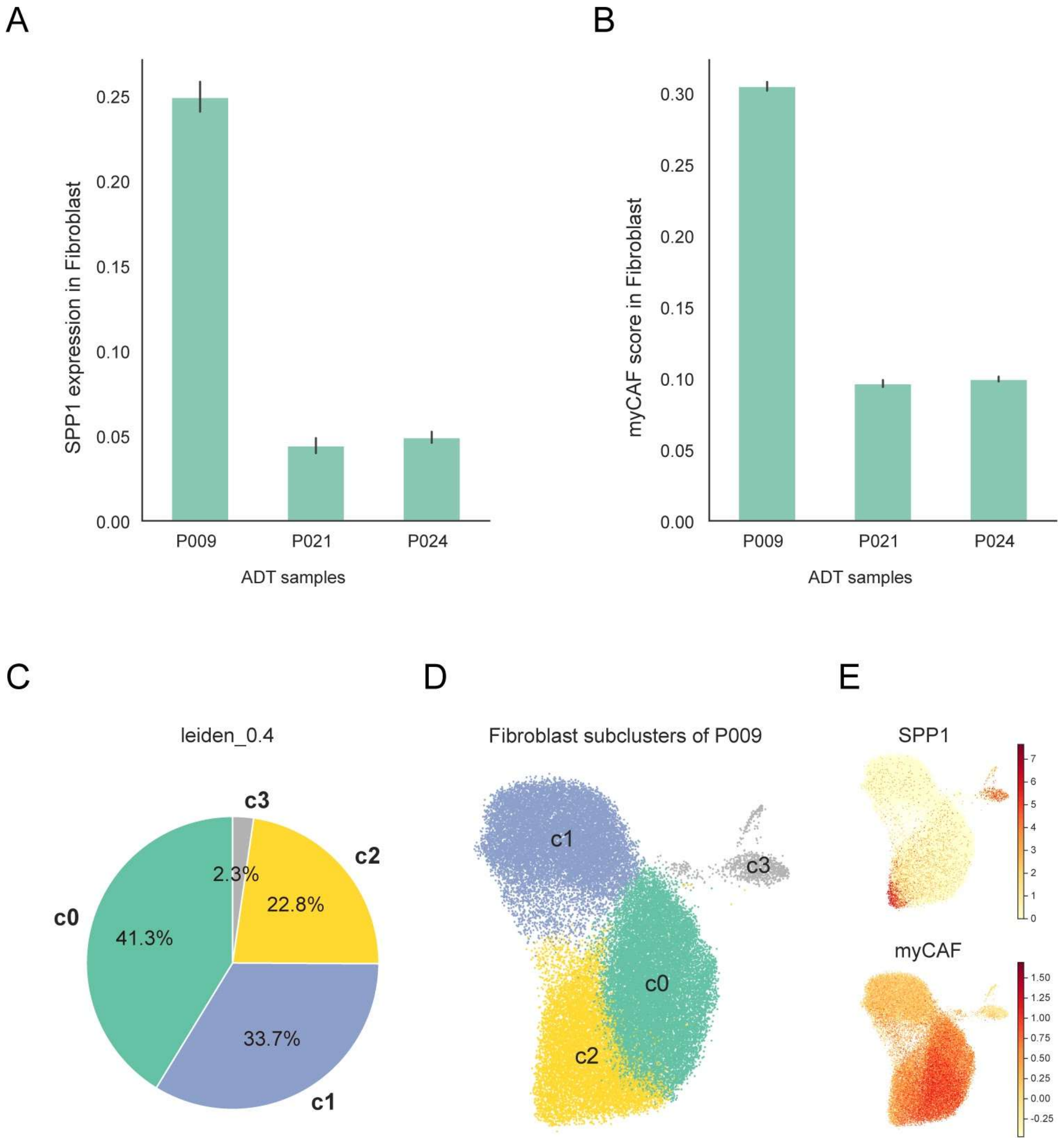


**Supplementary Fig. 4 | SPP1-associated fibroblast subpopulation analysis.** (**A**) Bar plots showing the mean SPP1 expression of PCa cells from different ADT samples; error bars indicate 95% confidence intervals. (**B**) Bar plots showing the mean myCAF scores of PCa cells in different ADT samples; error bars indicate 95% confidence intervals. (**C**) Pie chart showing the proportional composition of four fibroblast subclusters identified in sample P009. (**D**) UMAP visualization of fibroblasts from sample P009, colored by the four subclusters. (**E**) UMAP visualization of fibroblasts from sample P009 colored by SPP1 expression and myCAF activity score.

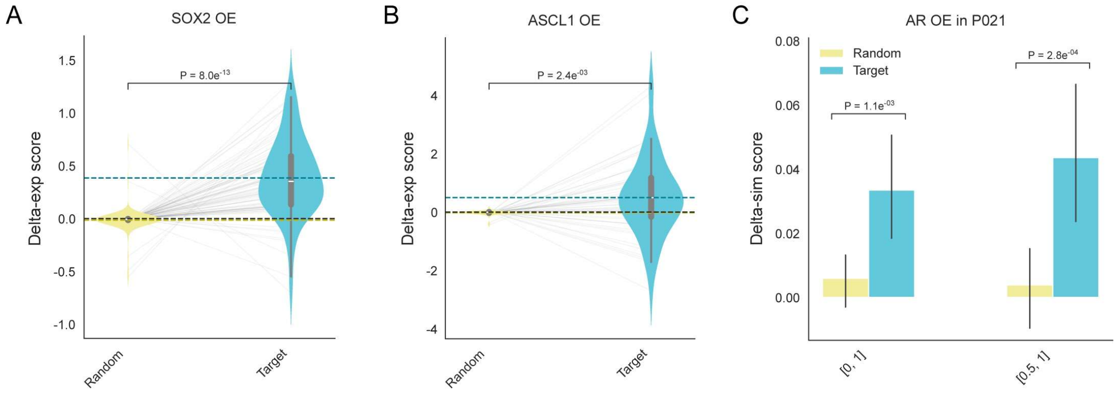


**Supplementary Fig. 5 | Identification of TFs related to NEPC by area ISP score.** (**A-B**) Violin plots showing the overall delta-exp scores for NEPC markers (CHGA, SYP, ENO2, and NCAM1) under target perturbations (**A**: SOX2 OE, **B**: ASCL1 OE) compared with random perturbations, as predicted by TMEformer. Statistical significance was assessed using a one-sided Wilcoxon rank-sum test. Dashed lines indicate group means (blue for Target; yellow for Random) and the zero baseline (black). (**C**), Bar plots showing the mean delta-sim scores of AR OE relative to random perturbations in the P021 sample; error bars indicate 95% confidence intervals. All PCa cells expressing AR are defined as [0,1], whereas low-AR cells ([0.5,1]) correspond to cells in which AR ranks in the bottom 50% of expressed genes.

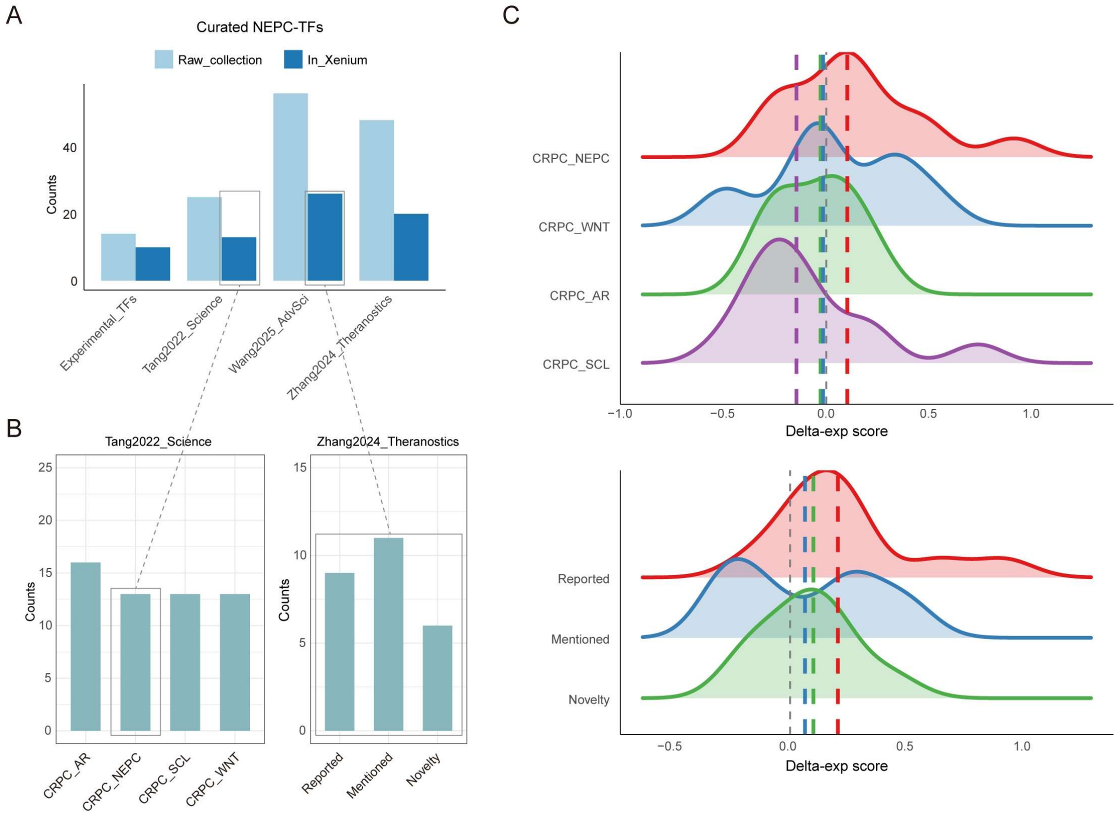


**Supplementary Fig. 6 | Collection and evaluation of NEPC-related TFs.** (**A**), Distribution of NEPC-related TFs curated from four resources, stratified by overlap with Xenium panel genes.

(**B**), Left: four CRPC subtype-associated TFs from Tange2022_Science. Right: three levels of NEPC-related TFs stratified by literature support, from Zhang2024_Theranostics. (**C**), Density plots of overall delta-exp area scores for NEPC markers (CHGA, SYP, ENO2, and NCAM1) among CRPC subtypes from Tange2022_Science set (Top) or literature support level from Zhang2024_Theranostics set (Bottom). Dotted lines indicate group medians.

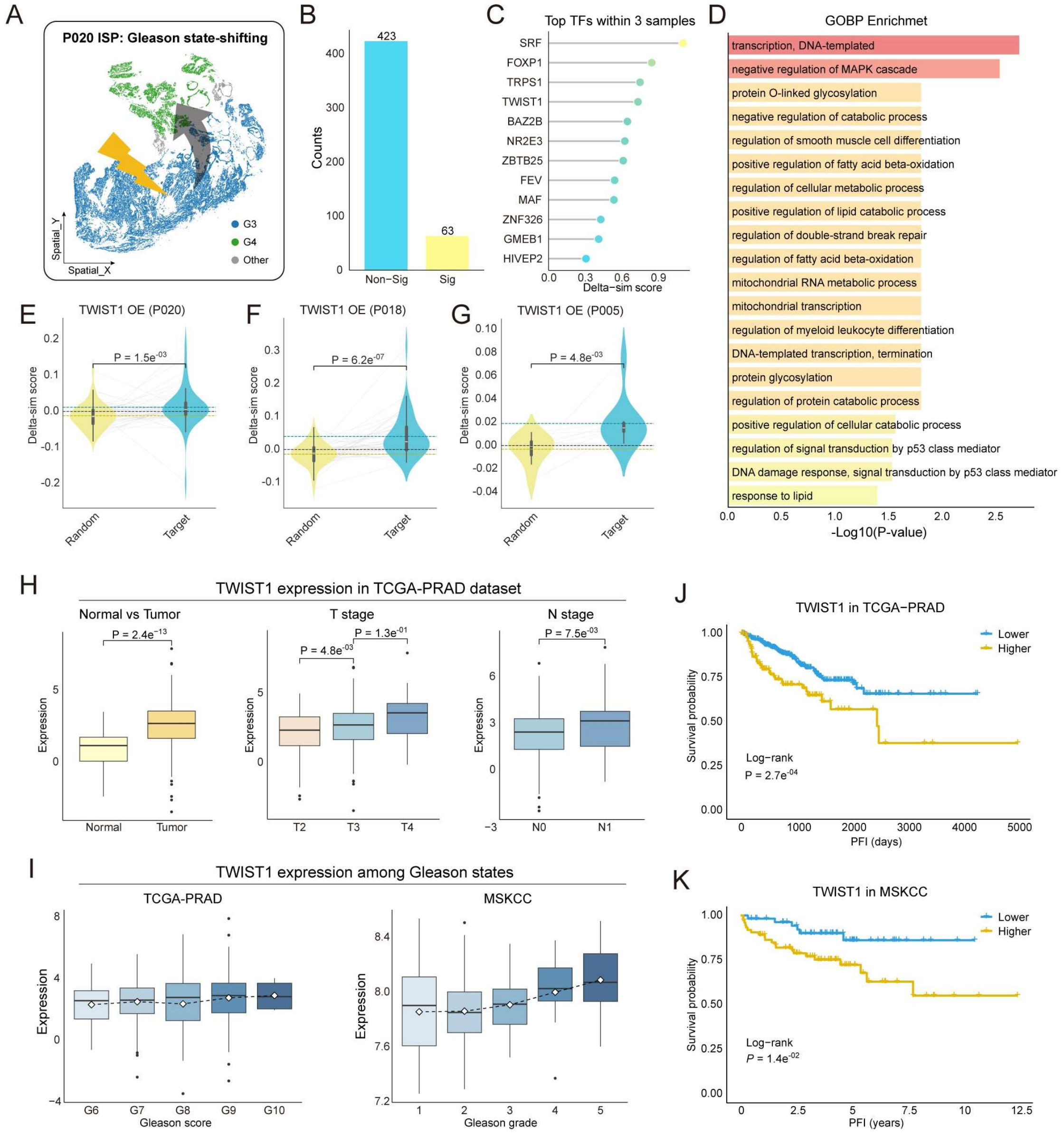


**Supplementary Fig. 7 | Identification of TFs promoting Gleason state transition in PCa.** (**A**), Schematic of embedding similarity–based ISP, evaluating whether perturbations in Gleason 3 (G3) region PCa cells increase their embedding similarity to Gleason 4 (G4) region PCa cells in P020 sample. (**B**), Bar plots showing the numbers of TFs with non-significant or significant effects toward G4 state, as predicted by TMEformer. (**C**), Mean normalized delta-sim scores of prioritized TFs with significant promoting effects toward G4 state across samples. (**D**), GOBP pathway enrichment analysis for significant TFs as reported in (**B**). (**E-G**), Violin plots showing the delta-sim scores toward the G4 state of TWIST1 OE compared with random perturbations in P020 (**E**), P018 (**F**), P005 (**G**) samples. (**H**), Boxplots comparing TWIST1 expression grouped by tissue types, T stages and N stages in TCGA-PRAD dataset. Statistical significance was assessed using Wilcoxon rank-sum tests. (**I**), Box plots comparing TWIST1 expression across Gleason states in the TCGA-PRAD and MSKCC datasets. Diamonds indicate group means. (**J**-**K**), Kaplan–Meier analyses of progression-free interval (PFI) stratified by TWIST1 expression in the TCGA-PRAD (**J**) and MSKCC (**K**) datasets using dataset-specific optimal cutoffs.

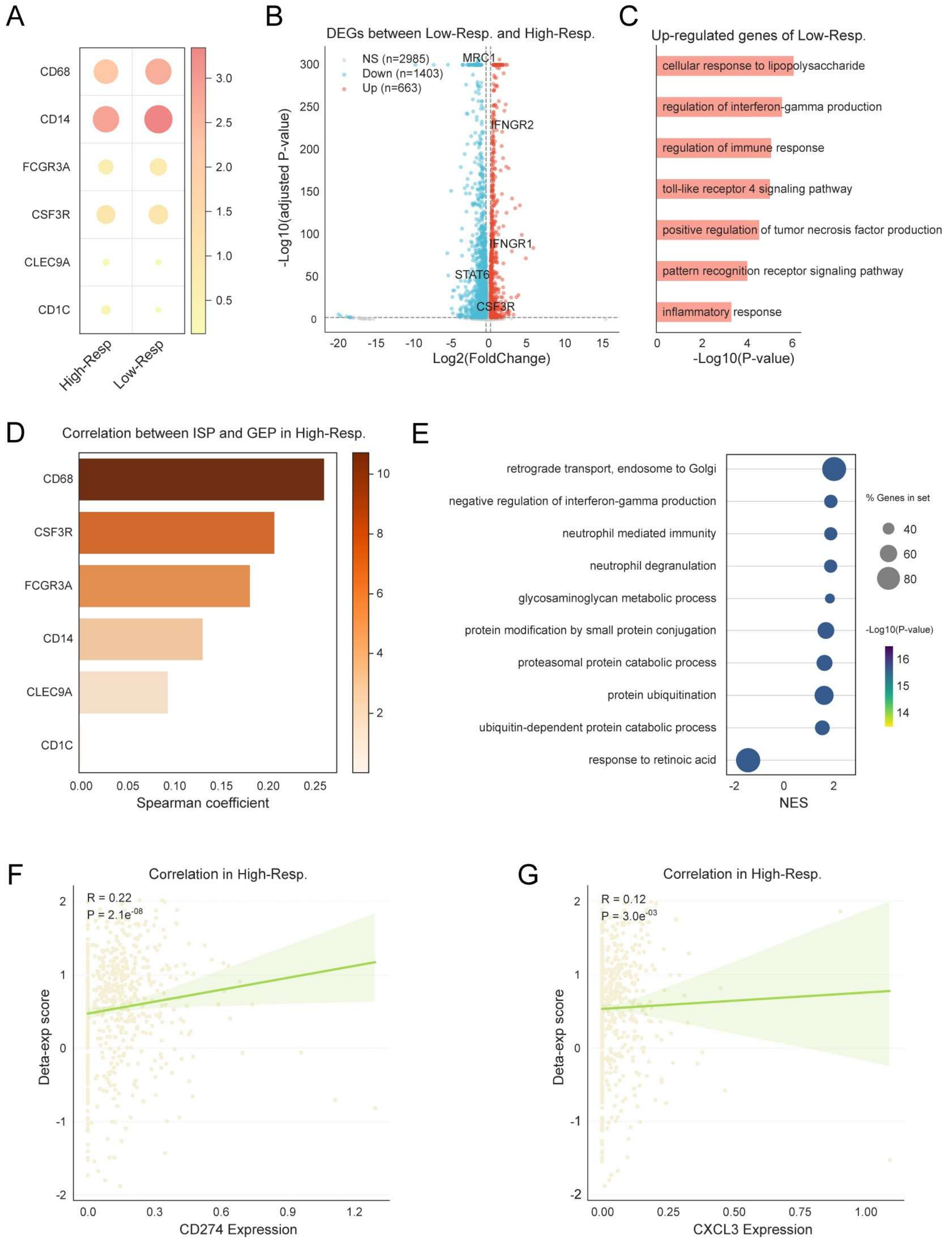


**Supplementary Fig. 8 | Characterization of Low-Resp and High-Resp grouped by perturbation effect of myeloid expansion.** (**A**) Dot plot comparing gene expression in myeloid cells between High-Resp and Low-Resp. Color indicates mean expression, and size indicates the percentage of expressing cells. (**B**) Volcano plot showing the differential expression genes (DEGs) in myeloid cells from Low-Resp versus High-Resp. (**C**) GOBP pathway enrichment analysis for up-regulated genes in Low-Resp. (**D**) Spearman correlation between ISP scores and myeloid subtype marker expression in High-Resp. (**E**), Enrichment analyses of GOBP pathways in genes ranked by correlation with ISP scores in High-Resp group. (**F**-**G**), Spearman correlation between ISP scores and CD274 (**F**) or CXCL3 (**G**) expression in myeloid cells of High-Resp group.

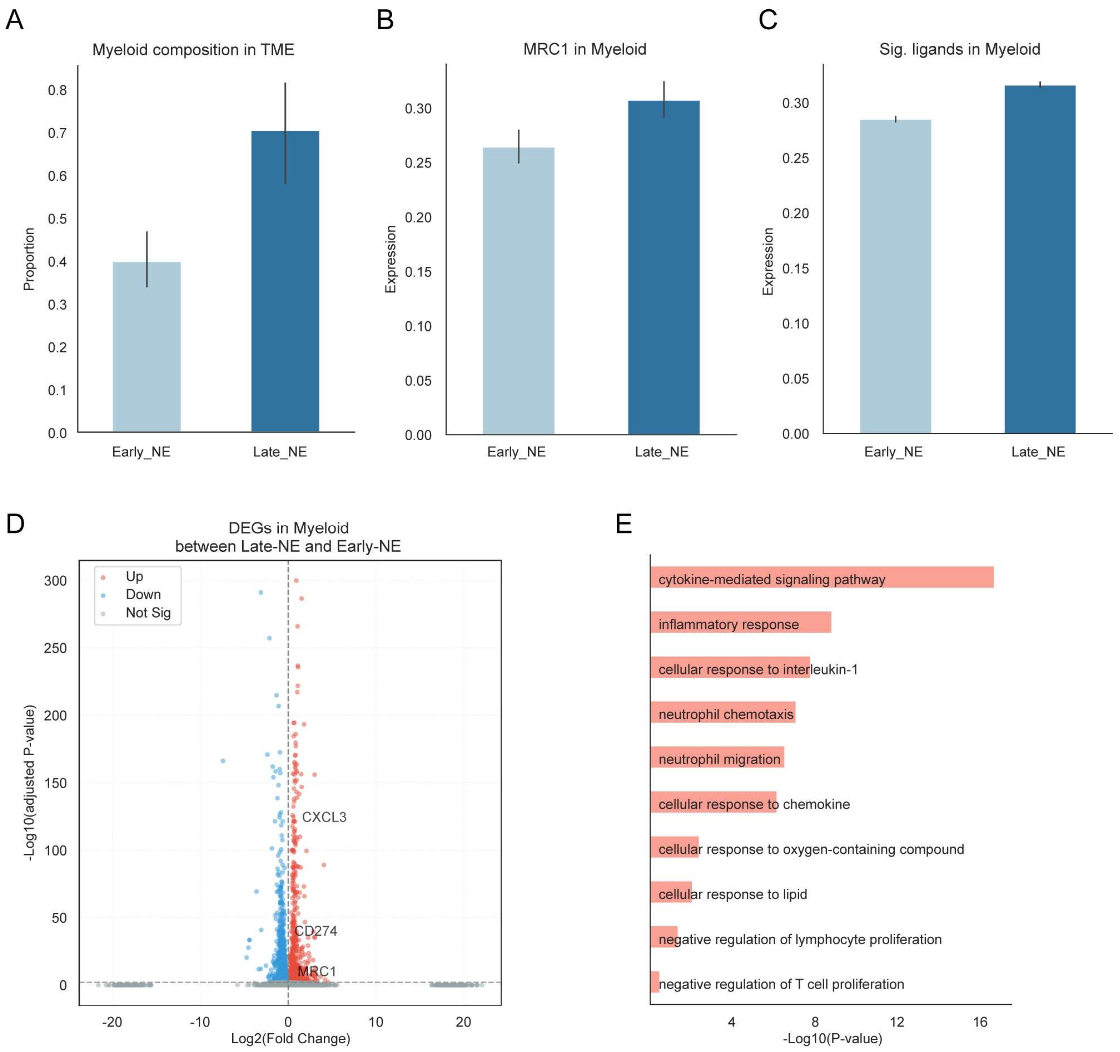


**Supplementary Fig. 9 | scRNA-seq analysis of myeloid cells between stages of NEPC modeling mouse.** (**A**) Bar plots showing mean myeloid proportions between Early_NE and Late_NE stages. (**B**) Bar plots showing mean MRC1 expression in myeloid between Early_NE and Late_NE stages. (**C**) Bar plots showing mean expression scores of ligands with significant ISP effects in myeloid between Early_NE and Late_NE stages. (**D**) Volcano plot showing the differential expression genes (DEGs) in myeloid cells from Late-NE versus Early-NE stages. (**E**) GOBP pathway enrichment analysis for up-regulated genes in myeloid cells of Late-NE stage.

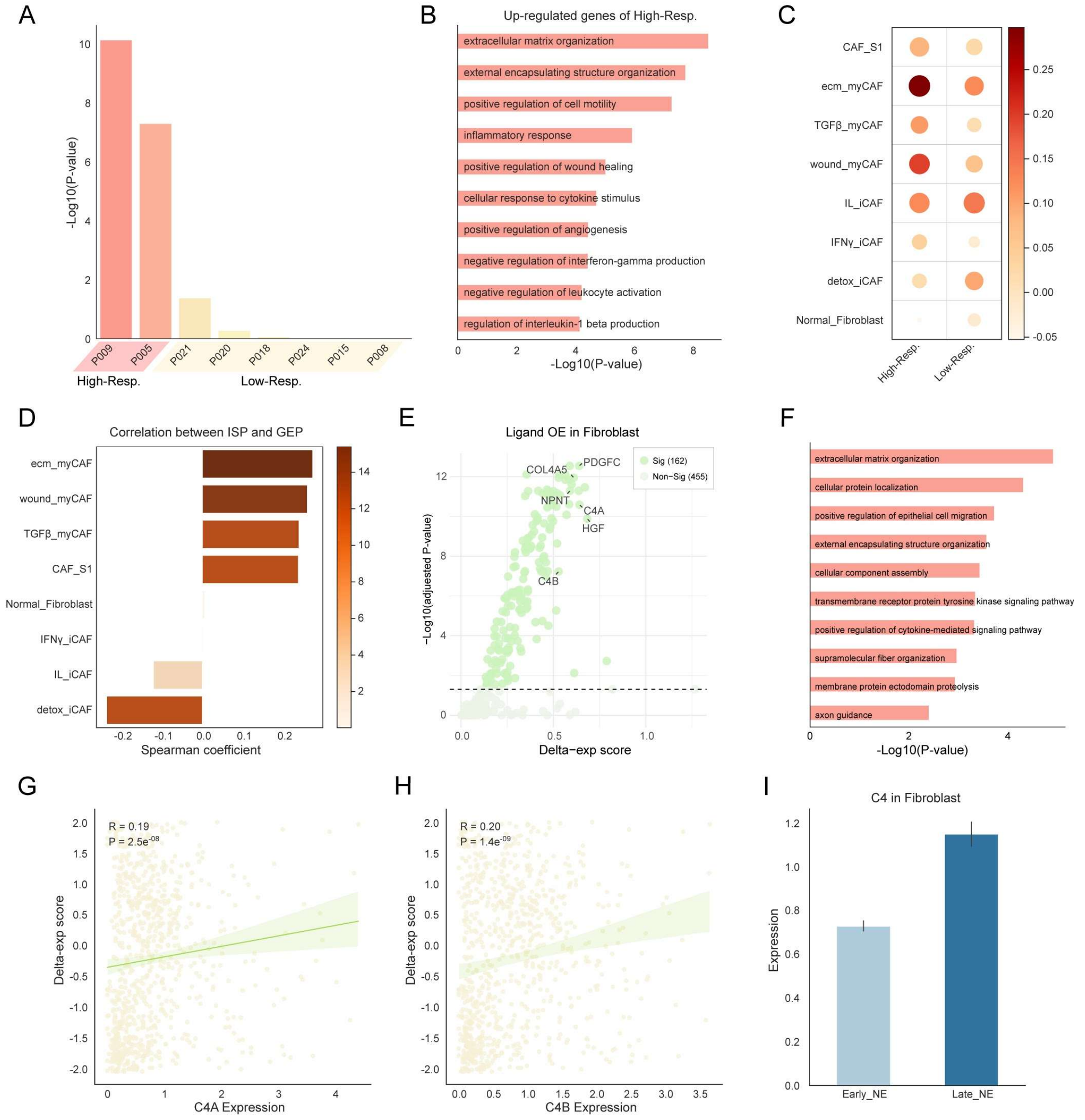


**Supplementary Fig. 10 | Fibroblast expansion and ligand perturbation analyses that drive NEPC expression.** (**A**) Bar plots show the significance of expanding fibroblasts in each sample. Samples with significant ISP effects on expanding myeloid cells are classified as High-Resp, and the remaining samples as Low-Resp. (**B**) GOBP pathway enrichment analysis for up-regulated genes in High-Resp. (**C**) Dot plot comparing gene expression in fibroblasts between High-Resp and Low-Resp. Color indicates mean expression, and size indicates the percentage of expressing cells. (**D**) Spearman correlation between ISP scores and fibroblasts-related signature expression scores. (**E**) Scatter plots showing the OE effect of ligands in fibroblasts on NEPC markers, with the x-axis indicating mean ISP score and the y-axis indicating significance relative to random perturbations. Ligands with mean ISP scores below 0 are omitted. (**F**) GOBP pathway enrichment analysis for ligands with significant ISP effects. (**G**-**H**) Spearman correlation between ISP scores and C4A (**G**) or C4B (**H**) expression in fibroblasts. (**I**) Bar plots showing mean C4 expression in fibroblasts between Early_NE and Late_NE stages of NEPC modeling mouse.

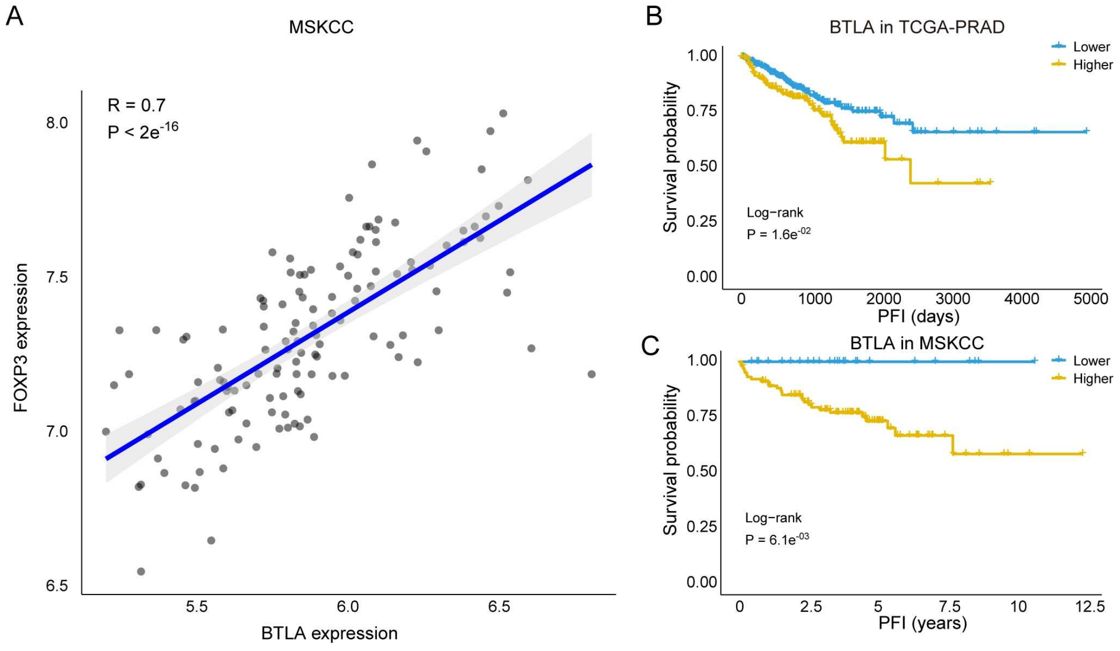


**Supplementary Fig. 11 | Validation of BTLA in MSKCC cohort**. (**A**) Spearman correlation between BTLA and FOXP3 expression in MSKCC cohort. (**B-C**) Kaplan–Meier survival analyses for progression-free interval (PFI) in TCGA-PRAD (**B**) and MSKCC (**C**) cohorts stratified by BTLA expression, using the optimal cutoff.

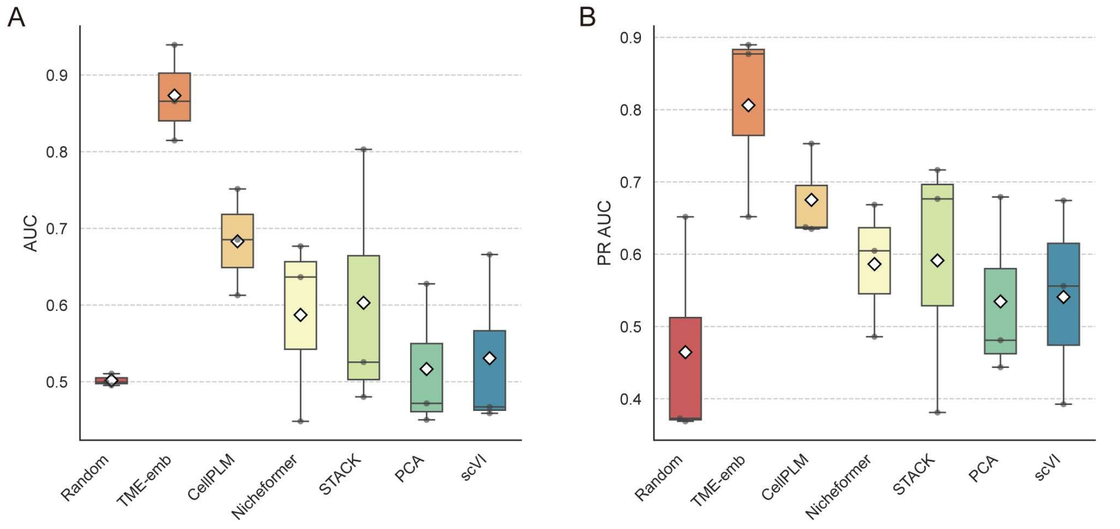


**Supplementary Fig. 12 | Validation of Gleason classifiers based on different cell embeddings.** Boxplots showing the AUC (**A**) and PR AUC (**B**) of model cross validation based on different cell embeddings. Each data point represents the performance on the remaining samples for a model trained on a single sample, with diamonds indicating the mean values

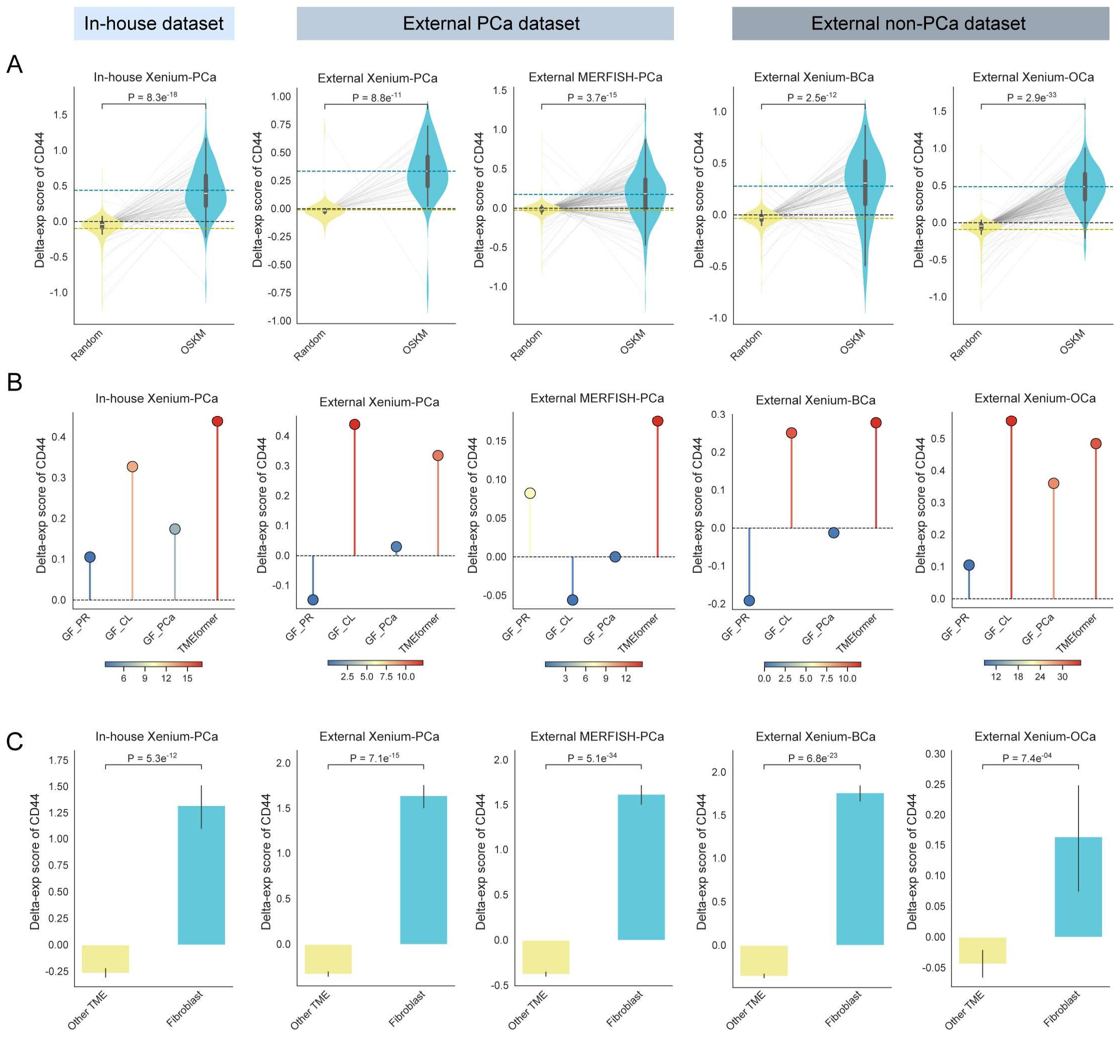


**Supplementary Fig. 13 | Perturbation effects on CD44 expression across spatial transcriptomics datasets.** (**A**), Violin plots showing delta-exp scores for CD44 under OSKM OE compared with random perturbations across in-house and external spatial transcriptomics datasets, as predicted by TMEformer. Statistical significance was assessed using a one-sided Wilcoxon rank-sum test. Dashed lines indicate group means (blue for Target; yellow for Random) and the zero baseline (black). (**B**), Dot plots showing the mean delta-exp scores for CD44 under OSKM OE across datasets. Color denotes statistical significance as −log10(P-value) relative to random perturbations using the same statistical framework as in (**A**). (**C**), Bar plots comparing delta-exp scores for CD44 following expansion of fibroblasts versus other TME cell populations. Statistical significance was assessed using a one-sided Wilcoxon rank-sum test.
